# Elusive *Plasmodium* Species Complete the Human Malaria Genome Set

**DOI:** 10.1101/052696

**Authors:** Gavin G. Rutledge, Ulrike Böehme, Mandy Sanders, Adam J. Reid, Oumou Maiga-Ascofare, Abdoulaye A. Djimdé, Tobias O. Apinjoh, Lucas Amenga-Etego, Magnus Manske, John W. Barnwell, François Renaud, Benjamin Ollomo, Franck Prugnolle, Nicholas M. Anstey, Sarah Auburn, Ric N. Price, James S. McCarthy, Dominic P. Kwiatkowski, Chris I. Newbold, Matthew Berriman, Thomas D. Otto

**Affiliations:** Malaria Programme, Wellcome Trust Sanger Institute, Hinxton, Cambridge, United Kingdom; Malaria Research and Training Center, University of Science, Techniques, and Technologies of Bamako, Bamako, Mali; Benhard-Nocht Institute for Tropical Medicine, Hamburg, Germany; University of Buea, Buea, Cameroon; Navrongo Health Research Centre, Navrongo, Ghana; Centers for Disease Control and Prevention, Atlanta, Georgia, United States; Laboratoire MIVEGEC (UM1-CNRS-IRD), Montpellier, France; Centre International de Recherches médicales de Franceville, Gabon; Global and Tropical Health Division, Menzies School of Health Research and Charles Darwin University, Darwin, NT, Australia; Centre for Tropical Medicine and Global Health, Nuffield Department of Clinical Medicine, University of Oxford, UK; Clinical Tropical Medicine Laboratory, QIMR Berghofer Medical Research Institute, University of Queensland, Brisbane, Australia; Wellcome Trust Centre for Human Genetics, University of Oxford, Oxford, United Kingdom; Weatherall Institute of Molecular Medicine, University of Oxford, John Radcliffe Hospital, Oxford, United Kingdom

## Abstract

Despite the huge international endeavor to understand the genomic basis of malaria biology, there remains a lack of information about two human-infective species: *Plasmodium malariae* and *P. ovale*. The former is prevalent across all malaria endemic regions and able to recrudesce decades after the initial infection. The latter is a dormant stage hypnozoite-forming species, similar to *P. vivax*. Here we present the newly assembled reference genomes of both species, thereby completing the set of all human-infective *Plasmodium* species. We show that the *P. malariae* genome is markedly different to other *Plasmodium* genomes and relate this to its unique biology. Using additional draft genome assemblies, we confirm that *P. ovale* consists of two cryptic species that may have diverged millions of years ago. These genome sequences provide a useful resource to study the genetic basis of human-infectivity in *Plasmodium* species.

## Introduction

All of the human malaria species were described in the early 20^th^ Century, with *Plasmodium malariae* and *P. ovale* being recognized as distinct species from *P. falciparum, P. vivax*, and *P. knowlesi*^*1*^. Reference genomes have now been published for the latter three^*2*^^−^^*4*^, with the extent of human infections caused by *P. knowlesi* having only been recognized decades after initial discovery^5^. Analysis of these reference genomes has revealed the basis of key biological processes, including virulence^6^, invasion^7^, and antigenic variation^8^. Despite the huge international endeavor to understand the genomic basis of malaria biology, almost nothing is known about the genetics of *P. malariae* and *P. ovale*.

Infections with these two organisms are frequently asymptomatic^9^ and have parasitaemia levels often below the level of detection of light microscopy^10^, thus making them difficult to study in human populations and potentially thwarting efforts to eliminate them and declare any region as ‘malaria free’^11^. This lack of knowledge is especially worrying because the two species are distributed widely across all malaria-endemic areas of the world^12^,^13^ (Figure 1a). Both species are frequent co-infections with the two common human pathogens, *P. falciparum* and *P. vivax*, and can be present in up to 5% of all clinical malaria cases^9^. This equates to roughly 30 million annual clinical cases. *P. malariae* infections can lead to lethal renal complications^14^ and can recrudesce decades later^15^, further increasing their socioeconomic costs.

**Figure 1.**
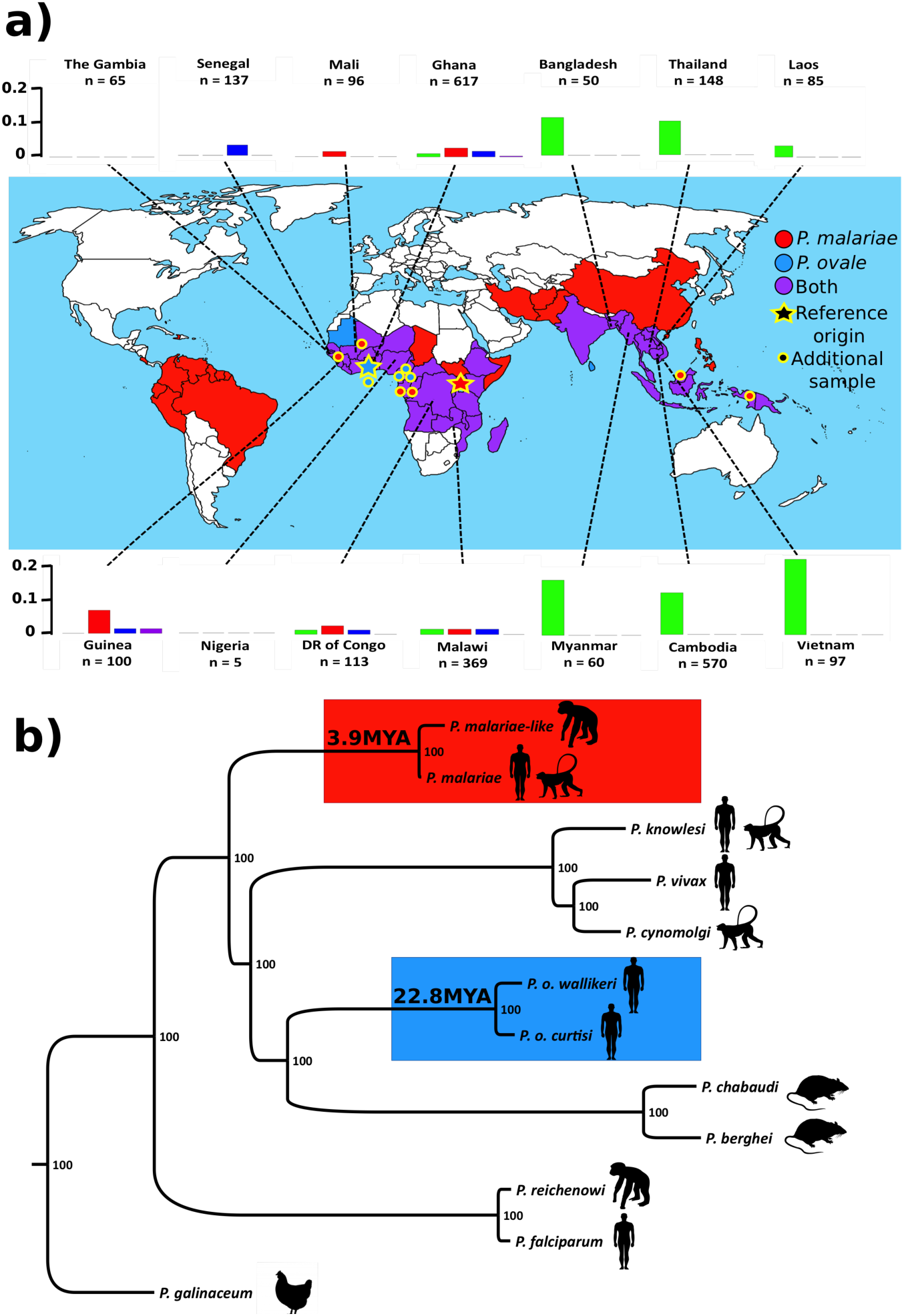
Prevalence and Phylogenetic Relationship of *P. malariae* and *P. ovale*. **a)**World map showing presence and absence of *P. malariae* (Red), *P. ovale* (Blue) or both (Purple) by country based on a literature review. Barplots show proportion of *P. falciparum* infections with co-infections of *P. malariae* (Red), *P. ovale* (Blue), *P. vivax* (Green), or two species (Purple) based on the Pf3K dataset. Stars indicate origin of sample used for reference genome assembly and points show additional samples used. **b)** Maximum likelihood phylogenetic tree of the *Plasmodium* genus, showing the *P. malariae* clade (Red) and the *P. ovale* clade (Blue) together with the divergence times of the species within those clades in millions of years ago (MYA). Silhouettes show infectivity of the different species.

Unraveling the mechanisms that enable *P. malariae* to persist in the host for decades is critical for a more general understanding of chronicity in malaria. The genome sequence of *P. ovale*, the other hypnozoite-forming species, will facilitate the search for conserved hypnozoite genes and will conclusively show whether *P. ovale* consists of two cryptic subspecies, as recently suggested^16^. Finally, the genetic basis of human-infectivity in malaria parasites can only be fully understood by having access to the genome sequences of all human-infective species.

Here we present the genome sequences of both these species, including the two recently described^16^ subspecies of *P. ovale (P. o. curtisi* and *P. o. wallikeri)*. We update the phylogeny of the *Plasmodium* genus using whole genome information, and describe novel genetic adaptations underlying their unique biology. Using whole genome sequencing of additional *P. malariae* (including two obtained from chimpanzees, referred to as *P. malariae-like)* and *P. ovale* samples, we describe the genetic variation present, as well as identify genes that are under selection. The data presented here provide the community with an essential foundation for further research efforts into these neglected species and into understanding the evolution of the *Plasmodium* genus as a whole.

## Results

### Plasmodium Co-Infections

Obtaining *P. malariae* and *P. ovale* DNA has historically been difficult due to the low level of parasitaemia in natural human infections. Using a novel method based on mitochondrial SNPs (Methods), we found *P. malariae* and *P. ovale* in approximately 2% of all *P. falciparum* clinical infections from the globally sampled Pf3K project (www.malariagen.net) (Figure 1a) (Supplementary Table 1), compared to 4% being co-infections with *P. vivax*. We also found a number of infections containing three species. These *P. malariae* and *P. ovale* co-infections are in addition to the larger number of mono-infections that they cause, which are frequently confounded by difficulties in confirming a species diagnosis. We used the two *P. ovale* co-infections with the highest number of sequencing reads to perform *de novo* genome assemblies (Supplementary Table 2).

### Genome Assemblies

A 33.6 megabase (Mb) reference genome of *P. malariae* was produced from clinically isolated parasites and sequenced using Pacific BioSciences long-read sequencing technology. The assembled sequence comprises 14 super-contigs representing the 14 chromosomes, with 6 chromosome ends extending into telomeres, and a further 47 unassigned subtelomeric contigs containing an additional 11 telomeric sequences (Table 1). Using existing Illumina sequence data from two patients primarily infected with *P. falciparum*, reads were extracted (Methods) and assembled into 33.5 Mb genomes for both *P. o. curtisi* and *P. o. wallikeri*, each assembly comprising fewer than 800 scaffolds. The genomes are significantly larger than previously sequenced *Plasmodium* species, and have isochore structures similar to those in *P. vivax*, with a higher AT content in the subtelomeres. In addition, a *P. malariae-like* genome was produced using Illumina sequencing from parasites isolated from a chimpanzee co-infected with *P. reichenowi*. The *P. malariae-like* genome was more fragmented than the other assemblies and its 23.7 Mb sequence misses most subtelomeric regions due to whole genome amplification prior to sequencing.

Most of the *P. malariae* genome is collinear with *P. vivax*, however we see two instances of large recombination breakpoints. The chromosomes syntenic to the *P. vivax* chromosomes 6 and 10 have recombined (Supplementary Figure 1a) and a large internal inversion has occurred on chromosome 5 (Supplementary Figure 1b), confirmed by mapping additional *P. malariae* samples back to the reference assembly.

**Table 1.**
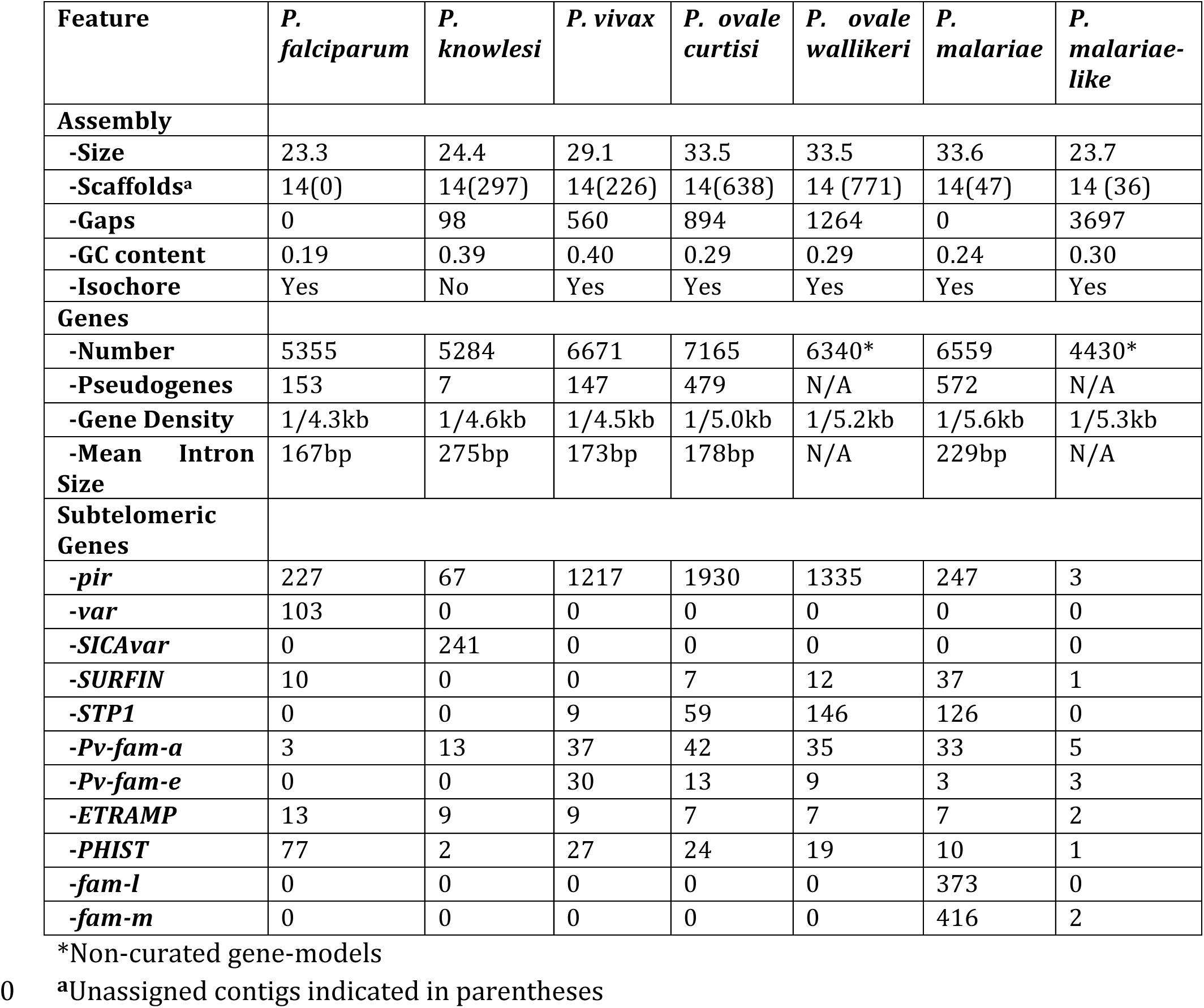
Comparison of Genome Features of all Human-Infective *Plasmodium* species and *P. malariae-like*

Across the four genomes, between 4,430 and 7,165 genes were identified using a combination of *ab initio* gene prediction and projection of genes from existing *Plasmodium* genome sequences. Manual curation was used to correct 2,516 and 2,424 genes for both the *P. malariae* and *P. o. curtisi* reference genomes respectively. A maximum likelihood tree was constructed using 1,000 conserved core genes that are present as single copies in 12 selected *Plasmodium* species (Figure 1b). The four newly assembled genomes do not cluster with any other *Plasmodium* species, but form two distinct and novel clades. Similar to a recent study using apicoplast data^17^, the two *P. ovale* species form a sister clade with the rodent malaria species, the latter being an ingroup to the ‘superfamily’ of primate-infective species in this tree. We also see that *P. malariae-like* has a longer branch length than *P. malariae*, which may be a reflection of the higher levels of diversity in *P. malariae-like* (Supplementary Figure 2a). This lack of diversity in *P. malariae* compared to the Chimpanzee species mirrors the situation of *P. falciparum* with *P. reichenowi^18^*.

We estimated the time of divergence for the four species using a Bayesian inference tool, G-PhoCS^19^. Absolute divergence time estimates are inherently uncertain due to mutation rate and generation time assumptions, and we therefore scaled these parameters to date the *P. falciparum* and *P. reichenowi* split using G-PhoCS to 4 million years ago (MYA), as previously published (3.0 − 5.5MYA)^20^. Assuming that the mutation rates and generation times are similar for *P. ovale* and *P. falciparum*, we find that the relative split of the two *P. ovale* species is about 5-times earlier than the split of *P. falciparum* and *P. reichenowi*. Using the same parameters as for the *Laverania* split, we thereby date the divergence of the two *P. ovale* subspecies to approximately 22.8MYA. This strongly supports the classification of *P. o. curtisi* and *P. o. wallikeri* as separate species rather than subspecies of *P. ovale*.

Using the same mutation rate and a longer generation time to account for the longer intra-erythrocytic cycle, we date the split of *P. malariae* from *P. malariae-like* to ~3.9MYA. This is similar to the estimated divergence of *P. falciparum* and *P. reichenowi*, suggesting a significant evolutionary event that promoted speciation in *Plasmodium* at that time. It has been suggested that a new world primate infective species termed *P. brasilianum* is identical to *P. malariae^21^*. To investigate this further using the new genome assemblies, we aligned the *P. brasilianum* merozoite surface protein 1 (MSP1)^22^ and ribosomal rRNA^21^ genes to both the *P. malariae* and *P. malariae-like* orthologous genes, showing that *P. brasilianum* is identical to *P. malariae*, but that *P. malariae-like* is indeed very different (Supplementary Figure 2b).

### Gene Changes

The greater number of genes in both *P. malariae* and *P. ovale* compared to existing *Plasmodium* genomes is mostly due to gene family expansions in the subtelomeres, such as *Plasmodium* interspersed repeat *(pir)* and *STP1* genes (Table 1). In addition, a large expansion of gamete antigen 25/27 (Pfg27) was identified in *P. malariae* with 22 tandemly duplicated copies including two pseudogenes on chromosome 14 (Supplementary Figure 3a). *P. vivax* and *P. falciparum* only have one and two copies respectively. Pfg27 is expressed highly during early gametocytogenesis^23^, and is essential for correct gametocyte development^24^. This gametocyte gene duplication may be an adaption by this species to ensure sexual reproduction in a setting of low level parasitaemia during infection.

In the *P. ovale* species, certain genes are also tandemly duplicated. Nine homologs (including two pseudogenes) of PVP01_1270800 are present in *P. o*. curtisi and 7 homologs are present in *P. o. wallikeri* (Supplementary Figure 3b). The *P. vivax* homolog is most highly expressed in sporozoites but has no known function^25^. The 3D structure of this gene, as predicted by I-TASSER^26^, appears to be similar to a nuclear pore complex (TM-Score > 0.4), suggesting a role in transport. This sporozoite change may be indicative of differences in liver-stage invasion or possibly hypnozoite formation.

Multiple genes have become pseudogenized in the two reference genomes compared to other human-infective *Plasmodium* species (Supplementary Table 3), including homologs of a multidrug efflux pump (PF3D7_0212800) which may suggest a higher susceptibility of these species to drug targeting. A phospho-fructo kinase, central to glycolysis^27^, is pseudogenized in both *P. ovale*, suggesting novel energy metabolism in these species. We also see genes that are pseudogenized in *P. o. wallikeri* but not *P. o. curtisi*, such as a serine-threonine protein kinase and a reticulocyte binding protein 1b (RBP1b), which is also pseudogenized in *P. malariae* as discussed below. One gene of specific interest that is pseudogenized in *P. o. wallikeri* but not in *P. o. curtisi* is a homolog of a cyclin in *P. falciparum* (PF3D7_1227500), an observation that may explain the different relapse times of the two *P. ovale* species^28^. The highest number of pseudogenes is seen in the *P. malariae* subtelomeres, where ~40% of the genes are pseudogenized in this species, indicating reduced selection pressure to cleanse the genome of these remnant genes.

*P. malariae* has a significantly longer intra-erythrocytic lifecycle compared to other human-infective *Plasmodium* species. All three *Plasmodium* cyclins^29^ are highly conserved in *P. malariae*, suggesting that the genetic cause may be elsewhere. A WD repeat-containing protein (WRAP73) is deleted in *P. malariae* but conserved across all other *Plasmodium* species. It is part of a large gene family known to be involved in a number of cellular processes, including cell cycle progression^30^. Knocking this gene out in other species may elucidate its importance in *Plasmodium* cell cycle progression.

Both *P. ovale* species are able to form hypnozoites, similar to *P. vivax3* and the simian-infective *P. cynomolgi^31^*. In searching for genes shared exclusively by these species, we identified 64 genes, of which two are of interest (Supplementary Table 4), as they do not belong to subtelomeric gene families. These include two conserved *Plasmodium* proteins, one of which has a low-level similarity to the *P. falciparum* ring-exported protein 4 gene. Both genes contain transmembrane domains and are expressed in *P. vivax* sporozoites^32^, making them interesting candidates to study experimentally.

### Subtelomeric Gene Families

The *Plasmodium* genus is characterized by species-specific subtelomeric gene family expansions, such as *var* genes in *P. falciparum*^*33*^ and *pir* genes in *P. yoelii*^34^. In *P. malariae* and *P. ovale*, where approximately 40% of the total genome size is subtelomeric, we also see large expansions of gene families that are species-specific (Figure 2a) (Table 1). The three largest gene clusters that we identified were in *P. malariae*. Of these, one cluster is composed of *STP1* and surface associated interspersed genes *(surfins). P. malariae* and *P. ovale* are the only human-infective species other than *P. falciparum*^*35*^ to contain *surfins* (Table 1), raising the possibility of studying this gene family using comparative genomics.

**Figure 2.**
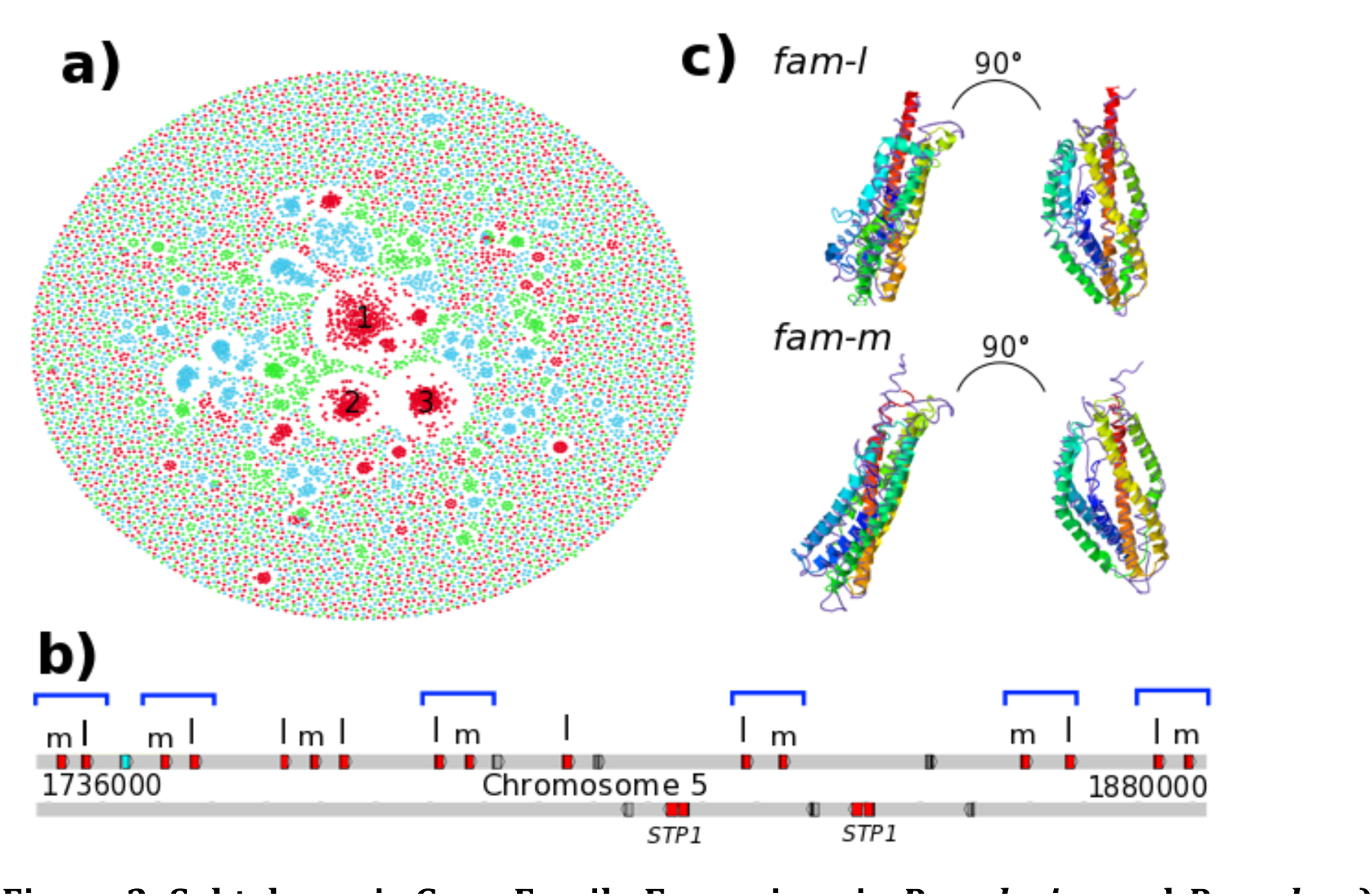
Subtelomeric Gene Family Expansions in *P. malahae* and *P. ovale*. **a)** Gene network based on sequence similarity of all genes in *P. malariae* (Red), *P. ovale* (Blue), and *P. vivax* (Green). Cluster 1 contains *fam-l* genes, Cluster 2 contains *fam-m* genes, and Cluster 3 contains *surfins* and *STP1* genes. **b)** Chromosome 5 subtelomeric localization of *fam-l* and *fam-m* genes in doublets (Blue brackets) on the telomere-facing strand. Also showing pseudogenes (Grey) and hypothetical gene (Blue). **c)** Predicted 3D-structure of *fam-l* (above) and *fam-m* (below) overlaid with the RH5 crystal structure (Purple). Left images show front of protein, right images show protein tilted to the right.

The other two large *P. malariae* clusters consist of two novel gene families, here termed *fam-l* and *fam-m*, consisting of 373 and 416 two-exon ~250 amino acid long genes respectively. We find two *fam-m* genes in *P. malariae-like*, which, despite the assembly lacking the majority of the subtelomeres, suggests that *P. malariae-like* also contains at least one of these novel families. The first exon of each *fam-l* and *fam-m* gene contains a signal peptide and a PEXEL motif-the signature in *P. falciparum* for export from the parasite into host erythrocytes^36^. In addition, the second exon contains two transmembrane domains flanking a hypervariable region. The remainder of the gene sequence is conserved between members of the same family and differentiates the two families from each other. These characteristics support the notion that the proteins coded for by these genes are exported from the parasite and may be targeted to the infected red blood cell surface and play a role in host-parasite interactions.

Ninety-three percent of *fam-l* and *fam-m* genes are on the same strand facing the telomeres (Figure 2b). This pattern, similar to *pir* genes in *P. yoelii^34^*, may be an adaptation to facilitate recombination between these genes. Uniquely, ~60% of these new genes are found in doublets of *a fam-l* and *a fam-m* (Figure 2b). Mirror tree analysis suggests that the pairs may be co-evolving over short periods of time (Supplementary Figure 4a), likely through being duplicated together, but that pairing may be disrupted by recombination over longer periods. We do not see any evidence of co-evolution between *pir* genes in close proximity of *fam-l* or *fam-m* genes (Supplementary Figure 4b), supporting the fact that this is not an artifact from their subtelomeric location. This suggests that *fam-l* and *fam-m* genes may encode proteins that dimerize when they are exported, a feature not previously seen among subtelomeric gene families.

Finally, we performed 3D structure prediction of both a *fam-l* and a *fam-m* gene using I-TASSER^26^. For both genes we got similar high-confidence (TM score > 0.5) 3D structures. These structures overlap the crystal structure of the *P. falciparum RH5* protein very well (TM score > 0.8), with 100% of the *RH5* structure covered even though they only have 10% sequence similarity (Figure 2c). *RH5* is a prime vaccine target in *P. falciparum* due to its essential binding to basigin during invasion^37^. The *RH5* kite-shaped fold is known to be present in RBP2a in *P. vivax*^38^, and may be a conserved structure necessary for the binding capabilities of all RH and RBP genes. This suggests that *fam-l* and *fam-m* genes may be involved in binding host receptors.

While neither *P. ovale* species has *fam-l* or *fam-m* genes, they both have large expansions of the *pir* gene family with 1,930 and 1,335 *pir* genes in *P. o. curtisi* and *P. o. wallikeri* respectively, while *P. malariae* only has 247 *pir* genes. This is the largest number of *pir* genes in any sequenced *Plasmodium* genome to date, explaining the large subtelomeres of this species. These *pir* genes form large species-specific clusters suggesting recent expansions (Figure 2a), but most closely resemble those in *P. vivax*. Many subfamilies of *pir* genes in *P. malariae* and *P. ovale* are shared with *P. vivax*, while almost none are shared with the rodent-infecting species (Supplementary Figure 5a). This suggests that *pir* genes are relatively well conserved between non-falciparum species infecting humans. Interestingly, all hypnozoite-forming species (Both *P. ovale, P. vivax*, and *P. cynomolgi)* contain over 1000 *pirs* each, significantly more than non-hypnozoite-forming *Plasmodium* species. Using additional draft genome assemblies for both *P. o. curtisi* and *P. o. wallikeri*, we show that the two species of *P. ovale* share significantly fewer *pir* genes inter-specifically than they do intra-specifically or intra-genomically (99% identity over 150 amino acids), further suggesting that the two species are not recombining with each other (Supplementary Figure 5b).

### Reticulocyte and Duffy Binding Proteins

RBP genes encode a merozoite surface protein family present across all *Plasmodium* species and known to be involved in red blood cell invasion and host specificity^39^. Compared to *P. vivax, P. malariae* has lost multiple RBPs including nearly all RBP2 genes and RBP1b, though it does have a functional RBP3. On the other hand, the two *P. ovale* species each have multiple full-length RBP2 genes (seven in *P. o. curtisi* and four in *P. o. wallikeri)* compared to three copies in *P. vivax* (Figure 3a). The two *P. ovale* species have very similar RBP2s, such as PocGH01_00019400 and PowCR01_00048600, a number of RBP2 pseudogenes in the two genomes match with a functional copy in the other genome (Supplementary Figure 6a). The RBP1b in *P. o. wallikeri* is less pseudogenized than the RBP1b in *P. malariae* and in *P. malariae-like* where we have a short fragment of the gene (Figure 3b). The specific mutation introducing a stop codon is conserved across the two *P. o. wallikeri* samples (Supplementary Figure 6b), indicating that RBP1b has become pseudogenized recently in *P. o. wallikeri*, or that the shortened form may be functional and has therefore been maintained under selection. It is interesting to note that the positioning of RBP1b and RBP1a is conserved across all these species, but not with the rodent malaria species.

**Figure 3.**
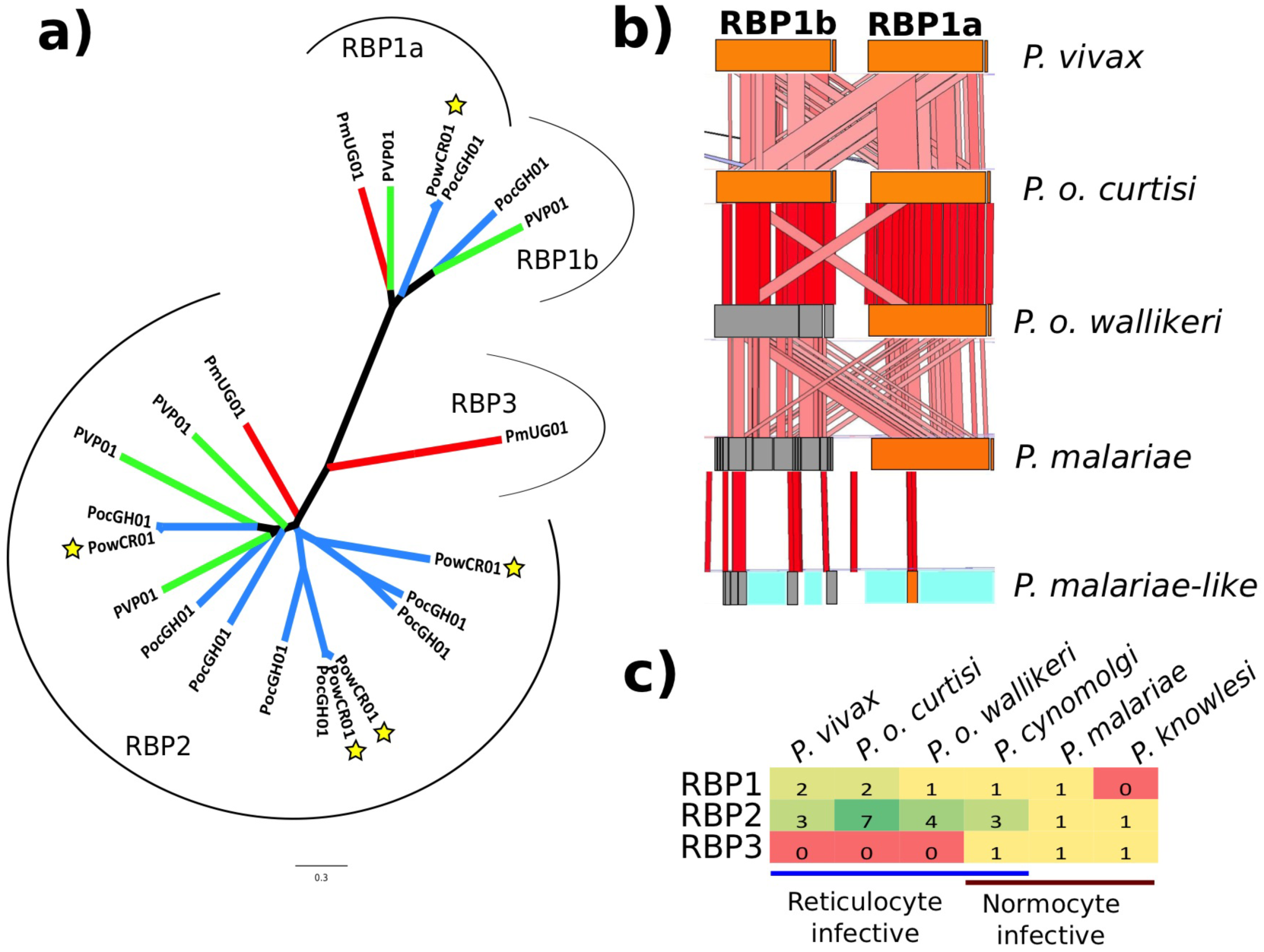
Reticulocyte Binding Protein Changes in *P. malariae* and *P. ovale*. **a)** Phylogenetic tree of all full-length functional RBPs in *P. malariae* (Red branches), *P. o. curtisi* (Blue branches without stars), *P. o. wallikeh* (Blue branches with stars), and *P. vivax* (Green branches). Brackets indicate the different subclasses of RBPs: RBP1a, RBP1b, RBP2, and RBP3. **b)** ACT67 view of functional (Orange) and pseudogenized (Grey) RBP1a and RBP1b in five species *(P. vivax, P. o. curtisi, P. o. wallikeh, P. malariae, P. malariae-like)*. Blue indicates assembly gaps. Red bars between species indicate level of sequence similarity, with darker colour indicating higher similarity. **c)** Number of RBP genes in each of the three RBP classes (RBP1, RBP2, RBP3) by species *(P. vivax, P. o. curtisi, P. o. wallikeri, P. cynomolgi, P. malariae, P. knowlesi)* grouped by erythrocyte invasion preference (reticulocyte versus normocyte).

RBP genes are thought to be involved specifically in reticulocyte invasion, which explains the gene loss in *P. malariae*, a species that preferentially invades normocytes^13^ (Figure 3c). Both *P. ovale* species exclusively invade reticulocytes^12^ and may have developed novel invasion pathways through the RBP2 expansion, similar to *P. vivax*. This supports a role for RBP2 gene expansions specifically in reticulocyte invasion. RBP3 genes seem to be pseudogenized in all reticulocyte-infective species, while they are fully functional in normocyte-infective species, suggesting a role in normocyte-invasion for RBP3.

Duffy binding proteins (DBPs) are also important for erythrocyte invasion^39^. *P. malariae* has one functional and one recently pseudogenized DBP, while both *P. ovale* have two functional copies. It is believed that *P. vivax* is incapable of infecting duffy-negative humans due to relying on its DBP binding the Duffy antigen, with recent studies showing duffy-negative infectivity in *P. vivax* strains containing a DBP duplication^40^. The fact that *P. malariae* and *P. ovale* are found throughout Africa (Figure 1a) suggests that they are capable of infecting duffy-negative individuals. It is therefore surprising that *P. malariae* only has one functional copy, implying that one copy is sufficient for duffy-negative infectivity in this species.

### Differential Selection Pressures

Using four additional *P. malariae* samples, two additional *P. o. curtisi* samples and two *P. malariae-like* and *P. o. wallikeri* samples each (Supplementary Table 2), we investigated differences in selection pressures between two species that diverged based on host differences *(P. malariae* and *P. malariae-like)*, and two species that supposedly diverged within the same host *(P. o. curtisi* and *P. o. wallikeri)*. Using GATK UnifiedGenotyper^41^, we called a total of 981,486 raw SNPs in *P. malariae* and 2,458,473 raw SNPs in *P. ovale*. Excluding subtelomeric regions, the pairwise nucleotide diversity between the different *P. malariae* samples is 4.7 × 10^−4^ and for the *P. o. curtisi* samples it is 3.8 × 10^−4^, which is significantly lower than similar estimates for *P. falciparum*^*42*^ and *P. vivax*^*43*^. Following SNP filtering (Methods), we retained on 230,881 SNPs in *P. malariae* with an average of 8,295 SNPs between the reference genome and the different *P. malariae* samples and with 150,832 SNPs on average with *P. malariae-like* (Supplementary Table 5). In *P. ovale* we retained 1,462,486 SNPs, of which 37,897 SNPs were different on average between *P. o. curtisi* samples and 1,412,799 were different on average between *P. o. curtisi* and *P. o. wallikeri* (Supplementary Table 6).

We calculated a number of selection measures for every core gene with 5 or more nucleotide substitutions (2,192 genes in *P. malariae*, 4,579 genes in *P. o. curtisi)*, including the Hudson-Kreitman-Aguade ratio (HKAr)^44^, which is the ratio of interspecific nucleotide divergence to intraspecific polymorphisms *(ie*. diversifying selection), Ka/Ks^45^, to look for an enriched number of nonsynonymous differences compared to synonymous differences *(ie*. positive selection), and the McDonald Kreitman (MK) Skew^46^, a measure of maintained polymorphisms *(ie*. balancing selection). We find high levels of HKAr (HKAr > 0.15) in a large proportion of genes in *P. malariae*, (127/2,192, 5.8%), but not in *P. o. curtisi* (36/4,579, 0.8%) (2-sample test for equality of proportions, p < 0.001) (Figure 4) (Supplementary Table 7). We see more genes under significant balancing selection in *P. malariae* (9/2,192, 0.4%) than in *P. o. curtisi* (5/4,579, 0.1%) (p < 0.05). More genes are under positive selection in *P. malariae* (104/2,192, 4.7%) than in *P. ovale* (24/4,579, 0.5%) (p < 0.001). This suggests that *P. malariae* may be under more widespread or stronger selective pressure than *P. o. curtisi*.

**Figure 4.**
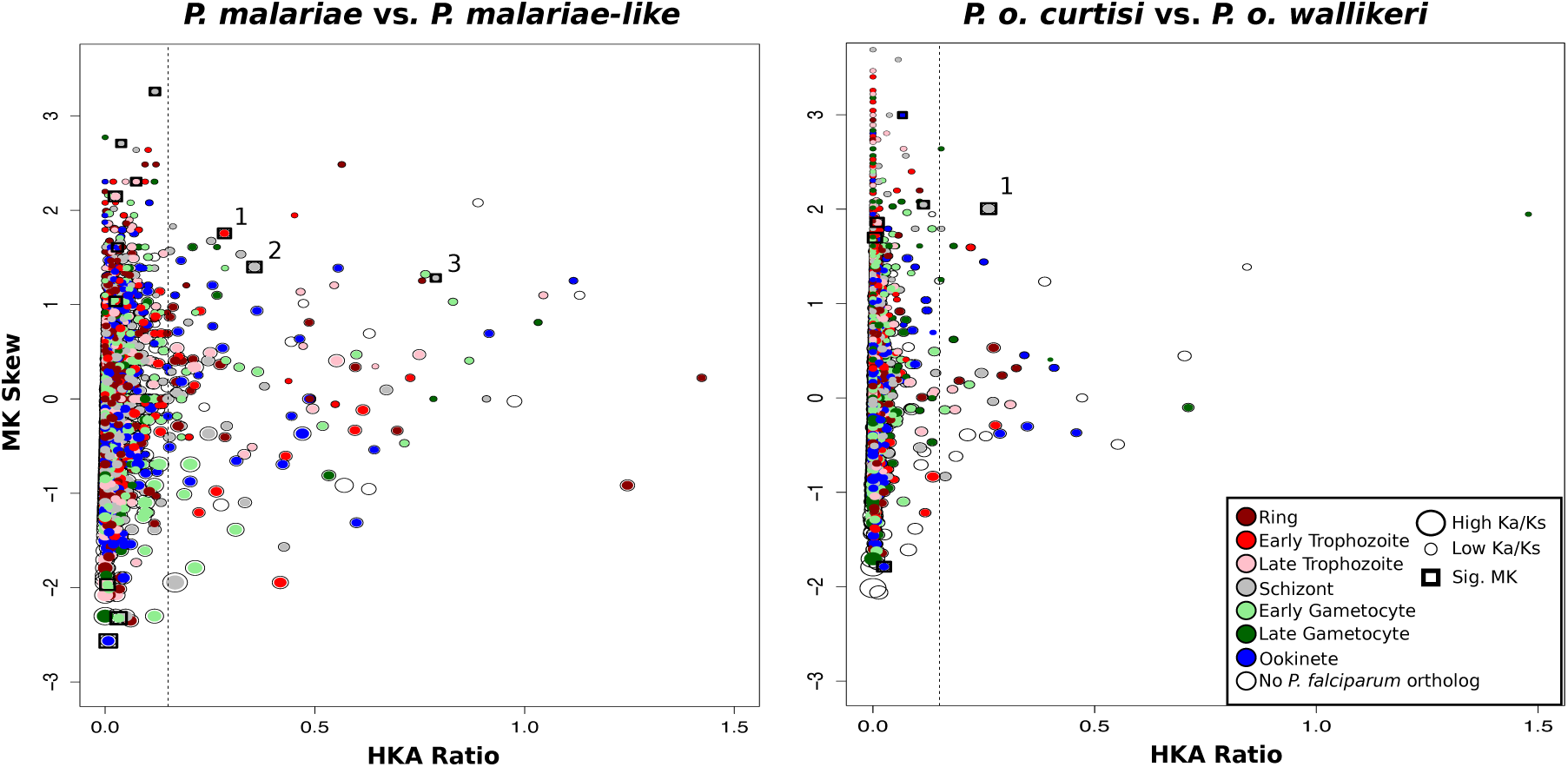
Differences in Gene-wide Selection Pressures between *P. malariae* and *P. ovale*. HKA ratio by MK Skew for both *P. malariae* versus *P. malariae-like* (left) and *P. o. curtisi* versus *P. o. wallikeri* (right) with the size of the point proportional to the Ka/Ks of that gene. Genes are coloured by their orthologs peak expression in *P. falciparum25* (Dark red for rings, red for early trophozoites, pink for late trophozoites, grey for schizonts, light green for early gametocytes, dark green for late gametocytes, blue for ookinetes, and blank for genes without a 1-1 ortholog in *P. falciparum)*. Genes with a significant MK skew are boxed in by a square. Genes right of the vertical dotted line have high HKAr. Genes with significant MK skews and high HKAr in *P. malariae* are: 1) rRNA methyl transferase 2) MSP1 3) formin-1, and for *P. o. curtisi:* 1) RBP2.

Looking at specific genes under selection, we see similar genes in the *P. malariae /P. malariae-like* test as in an earlier *P. falciparum/P. reichenowi* study^47^. This includes a large number of invasion genes with high HKAr values, such as MSP8 and MSP7, as well as significant MK skews for MSP1 and apical membrane antigen 1. For *P. malariae*, genes with high HKAr values besides invasion genes are associated with stages throughout the parasites lifecycle (Figure 4). However in *P. o. curtisi*, they are predominantly invasion and gametocyte genes, including among others a gametocyte associated protein and a mago nashi homolog protein (Figure 4), the latter potentially being involved in sex determination^48^. For *P. o. curtisi*, we also find a large number of genes with high Ka/Ks values that are gametocyte-associated, such as a sexual stage antigen s16. We therefore find that invasion genes tend to always be under strong selective pressure in *Plasmodium*, but that *P. malariae* and *P. o. curtisi* differ in terms of the other life cycle stages that are under selective pressure.

One of the genes with the highest Ka/Ks in the P*. malariae/P. malariae-like* comparison is RBP1a, which has 37 nonsynonymous fixed differences between the two species and only 6 synonymous fixed differences. The other two intact RBPs are much more highly conserved. Knowing that *P. malariae* also infects new world monkeys (where it is known as *P. brasilianum*)^21^, we might suppose that the receptor for RBP1a may be conserved between humans and new world monkeys, but not with chimpanzees. We identified 19 human genes coding for transmembrane-containing proteins that may act as potential RBP1a receptors (Methods), which includes a mucin-22 precursor and an aquaporin 12b precursor (Supplementary Table 8).

### Discussion

The high-quality genome sequences of *P. malariae* and *P. ovale* and their annotation presented here provide a rich new resource for comparative *Plasmodium* genomics. They provide a foundation for further studies into the biology of these two neglected malaria species, as well as new tools to explore genus level similarities and differences in infection. The genome sequences have revealed a number of genomic adaptations and possible consequences related to the success of these species sustaining low parasitaemia infections, including gametocyte gene expansions and an increase in genome size. The genome sequences suggest that the rodent-infective malaria species may be the result of an ancestral host switch from a primate-infective species and also conclusively show that *P. ovale* is a species complex, consisting of two highly diverged species, *P. o. curtisi* and *P. o. wallikeri*. The genome sequences reveal a novel type of subtelomeric gene family in *P. malariae* occurring in doublets and potentially having an *RH5-like* fold. Having access to a larger number of genome sequences also allows us to identify features such as the RBP2 gene expansion in reticulocyte invading *Plasmodium* species. Multi-sample analysis of the two species highlights differences in selection pressures between host-switching and within-host speciation, as well as the omnipresent selective pressure during red blood cell invasion. These genome sequences will now enable more comprehensive studies of human-infectivity in *Plasmodium* species.

## Methods

### Co-infection Mining

We aligned the *P. malariae* (AB354570) and *P. ovale* (AB354571) mitochondrial genome sequences against those of *P. falciparum*^*2*^, *P. vivax*^*3*^, and *P. knowlesi*^*4*^ using MUSCLE^49^. For each species, we identified three 15bp stretches within the *Cox1* gene that contained two or more species-specific SNPs. We searched for these 15bp species-specific barcodes within the sequencing reads of all 2,512 samples from the Pf3K global collection ( www.malariagen.net). Samples that contained at least two sequencing reads matching one or more of the 15bp barcodes for a specific species were considered to be positive for that species (Supplementary Table 1). We found good correspondence between the three different barcodes for each species, with over 80% of positive samples being positive for all three barcodes. We generated pseudo-barcodes by changing two randomly selected nucleotide bases at a time for 10 randomly selected 15bp region in the *P. vivax3* mitochondrial genome. We did not detect any positive hits using these pseudo-barcodes. As an additional negative control, we searched for *P. knowlesi* co-infections, but did not find any samples positive for this species. Two samples (PocGH01, PocGH02) had high numbers for all three *P. ovale* barcodes and were used for reference genome assembly and SNP calling respectively.

### Parasite Material

All *P. ovale* samples were obtained from symptomatic patients diagnosed with a *P. falciparum* infection. The two *P. o. curtisi* samples (PocGH01, PocGH02) identified through co-infection mining (see above), were from two patients testing positive on a CareStart^®^ (HRP2 based) rapid malaria diagnostic test (RDT) kit at the Navrongo War Memorial hospital, Ghana. Following consent obtainment, about 2-5mls of venous blood was obtained and then diluted with one volume of PBS. This was passed through CF11 cellulose powder columns to remove leucocytes prior to parasite DNA extraction.

The two *P. malariae-like* samples, PmlGA01 and PmlGA02, were extracted from Chimpanzee blood obtained during routine sanitary controls of animals living in a Gabonese sanctuary (Park of La Lékédi, Gabon). Blood collection was performed following international rules for animal health. Within six hours after collection, host white blood cell depletion was performed on fresh blood samples using the CF11 method^50^. After DNA extraction using the Qiagen blood and Tissue Kit and detection of *P. malariae* infections by *Cytb* PCR and sequencing^51^, the samples went through a whole genome amplification step^52^.

One *P. malariae* sample, PmGN01, collected from a patient with uncomplicated malaria in Faladje, Mali. Venous blood (2-5mL) was depleted of leukocytes within 6 hours of collection as previously described^53^. The study protocol was approved by the Ethics Committee of Faculty of Medicine and Odontomatology and Faculty of Pharmacy, Bamako, Mali.

Four samples of *P. malariae* were obtained from travellers returning to Australia with malaria. PmUG01 and PmID01 were sourced from patients returning from Uganda and Papua Indonesia respectively, who presented at the Royal Darwin Hospital, Darwin, with microscopy-positive *P. malariae* infection. PmMY01 was sourced from a patient presenting at the Queen Elizabeth Hospital, Sabah, Malaysia, with microscopy-positive *P. malariae* infection. Patient sample PmGN02 was collected from a patient who presented to Royal Brisbane and Womens Hospital in 2013 on return from Guinea.

Venous blood samples were subject to leukodepletion within 6 hours of collection. PmUG01 was leukodepleted using a commercial Plasmodipur filter (EuroProxima, The Netherlands); home-made cellulose-based filters were used for PmID01 and PmMY01, while PmGN02 was leukodepleted using an inline leukodepletion filter present in the venesection pack (Pall Leukotrap; WBT436CEA). DNA extraction was undertaken on filtered blood using commercial kits (QIAamp DNA Blood Midi kit, Qiagen Australia).

For samples PmUG01, PmID01 and PmMY01, ethical approval for the sample collection was obtained from the Human Research Ethics Committee of NT Department of Health and Families and Menzies School of Health Research (HREC-2010-1396 and HREC-2010-1431) and the Medical Research Ethics Committee, Ministry of Health Malaysia (NMRR-10-754-6684). For sample PmGN02, ethical approval was obtained from the Royal Brisbane and Womens Hospital Human Research Ethics Committee (HREC/10/QRBW/379) and the Human Research Ethics Committee of the Queensland Institute of Medical Research (p1478).

### Sample Preparation and Sequencing

One *P. malariae* sample, PmUG01, was selected for long read sequencing, using Pacific Biosciences (PacBio), due to its low host contamination and abundant DNA. Passing through a 25mm blunt-ended needle, 6ug of DNA was sheared to 20-25kb. SMRT bell template libraries were generated using the PacBio issued protocol (20kb Template Preparation using the BluePippin™ Size-Selection System). After a greater than 7kb size-selection using the BluePippin™ Size-Selection System (Sage Science, Beverly, MA), the library was sequenced using P6 polymerase and chemistry version 4 (P6/C4) in 20 SMRT cells (Supplementary Table 2).

The remaining isolates were sequenced with Illumina Standard libraries of 200-300bp fragments and amplification-free libraries of 400-600bp fragments were prepared^54^ and sequenced on the Illumina HiSeq 2000 v3 or v4 and the MiSeq v2 according to the manufacturers standard protocol (Supplementary Table 2). Raw sequence data was deposited in the European Nucleotide Archive (Supplementary Table 2).

### Genome Assembly

The PacBio sequenced *P. malariae* sample, PmUG01, was assembled using HGAP^55^ with an estimated genome size of 100Mb to account for the host contamination (~85% Human). The resulting assembly was corrected initially using Quiver^55^, followed by iCORN^56^. PmUG01 consisted of two haplotypes, with the majority haplotype being used for the iCORN56, and a coverage analysis was performed to remove duplicate contigs. Additional duplicated contigs were identified using a BLASTN^57^ search, with the shorter contigs being removed if they were fully contained within the longer contigs or merged with the longer contig if their contig ends overlapped. Host contamination was removed by manually filtering on GC, coverage, and BLASTN hits to the non-redundant nucleotide database^57^.

The Illumina based genome assemblies for *P. o. curtisi, P. o. wallikeri*, and *P. malariae-like* were performed using MaSURCA^58^ for samples PocGH01, PowCR01, and PmlGA01 respectively. To confirm that the assemblies were indeed *P. ovale*, we mapped existing *P. ovale* capillary reads to the assemblies (www.ncbi.nlm.nih.gov/Traces/trace.cgi?view=search). Prior to applying MaSURCA^58^, the samples were mapped to the *P. falciparum* 3D7 reference genome^2^ to remove contaminating reads. The draft assemblies were further improved by iterative uses of SSPACE^59^, GapFiller^60^ and IMAGE^61^. The resulting scaffolds were ordered using ABACAS^62^ against the *P. vivax* PVP01(href="http://www.genedb.org/Homepage/PvivaxP01) assembly (both *P. ovale)* or against the *P. malariae* PacBio assembly *(P. malariae-like)*. The assemblies were manually filtered on GC, coverage, and BLASTN hits to the non-redundant nucleotide database^57^. iCORN^56^ was used to correct frameshifts. Finally, contigs shorter than 1 kilobase (kb) were removed.

Using two more samples, PocGH02 and PowCR02, additional draft assemblies of both *P. ovale* species were produced using MaSURCA^58^ followed by RATT^63^ to transfer the gene models from the high-quality assemblies.

The genome sequences and annotation for both *P. malariae* and *P. ovale* can now be found on GeneDB at http://www.genedb.org/Homepage/Pmalariae and at http://www.genedb.org/Homepage/Povale.

### Gene Annotation

RATT^63^ was used to transfer gene models based on synteny conserved with other sequenced *Plasmodium* species *(P. falciparum*^2^, *P. vivax*^3^, *P. berghei*^34^, and *P. gallinaceum* (unpublished)). In addition, genes were predicted *ab initio* using AUGUSTUS^64^, trained on a geneset consisting of manually curated *P. malariae* and *P. ovale* genes respectively. Non-coding RNAs and tRNAs were identified using Rfam 12.0^65^. Gene models were then manually curated for both the *P. malariae* and *P. o. curtisi* reference genomes, using Artemis^66^ and the Artemis Comparison Tool (ACT)^67^. These tools were also used to manually identify deleted and disrupted genes (Supplementary Table 3).

### Phylogenetics

Following ortholog assignment using BLASTP^57^ and OrthoMCL^68^, amino acid sequences of 1000 core genes from 12 *Plasmodium* species *(P. galinaceum* (unpublished), *P. falciparum*^*2*^, *P. reichenowi*^*47*^, *P. knowlesi*^*4*^, *P. vivax*^*3*^, *P. cynomolgi*^31^, *P. chabaudi*^34^, *P. berghei*^34^, and the four assemblies produced in this study) were aligned using MUSCLE^49^. The alignments were cleaned using GBlock^69^ with default parameters to remove non-informative and gapped sites. The cleaned non-zero length alignments were then concatenated. This resulted in an alignment of 421,988 amino acid sites per species. The optimal substitution model for each gene partition was determined by running RAxML70 for each gene separately using all implemented substitution models. The substitution models that generated the tree with the highest likelihood were used for each gene partition. A maximum likelihood phylogenetic tree was constructed using RAxML^70^ with 100 bootstraps^71^ (Figure 1b). To confirm this tree, we utilized different phylogenetic tools including PhyloBayes^72^ and PhyML^73^, a number of different substitution models within RAxML, starting the tree search from the commonly accepted phylogenetic tree, and removing sites in the alignment which supported significantly different trees. All approaches yielded the final tree found in Figure 1b with highest likelihood. Figtree was used to colour the tree (http://tree.bio.ed.ac.uk/software/figtree/).

A phylogenetic tree of four *P. malariae* (PmID01, PmGN01, PmGN02, PmMY01) and all *P. malariae-like* samples (PmlGA01, PmlGA02) was generated using PhyML^73^ based on all *P. malariae* genes. For each sample, the raw SNPs as called using the SNP pipeline (see below), were mapped onto all genes to morph them into sample specific gene copies using BCFtools^74^. Amino acids for all genes were concatenated and cleaned using GBlocks^69^.

### Divergence Dating

Species divergence times were estimated using the Bayesian inference tool G-PhoCS^19^, a software which uses thousands of unlinked neutrally evolving loci and a given phylogeny to estimate demographic parameters. One additional sample per assembly (PmGN01 for *P. malariae*, PocGH02 for *P. o. curtisi*, PowCR02 for *P. o. wallikeri*, and PmlGA02 for *P. malariae-like)* was used to morph the respective assembly using iCORN^56^. Regions in the genomes without mapping were masked, as iCORN^56^ would not have morphed them. Unassigned contigs and subtelomeric regions were removed for this analysis due to the difficulty of alignment. Repetitive regions in the chromosomes of the four assemblies and the four morphed samples were masked using Dustmasker^75^ and then the chromosomes were aligned using FSA^76^. The *P. o. wallikeri* and the *P. o. curtisi* chromosomes were aligned against each other, as were the *P. malariae* and *P. malariae-like* chromosomes. The alignments were split into 1kb loci, removing those that contained gaps, masked regions, and coding regions to conform with the neutral loci assumption of G-PhoCS^19^. G-PhoCS^19^ was run for one million MCMC-iterations with a sample-skip of 1,000 and a burn-in of 10,000 for each of the two species pairs. Follow-up analyses using Tracer (http://beast.bio.ed.ac.uk/Tracer) confirmed that this was sufficient for convergence of the MCMC chain in all cases. In the model, we assumed a variable mutation rate across loci and allowed for on-going gene flow between the populations. The tau values obtained from this were 0.0049 for *P. malariae* and 0.0434 for *P. ovale*.

The tau values were used to calculate the date of the split, using the formula (tau x G)/mu, where G is the generation time in years and mu is the mutation rate. Following optimization of the *P. falciparum/P. reichenowi* split to 4 million years ago (unpublished), as estimated previously^20^, we assumed a mutation rate of 3.8 × 10^−10^ SNPs/site/lifecycle^77^ and a generation time of 65 days^78^. For *P. malariae*, a generation time of 100 days was used due to the longer intra-erythrocytic cycle.

### 3D Structure Prediction

The I-TASSER^26^ Version 4.4 online web server^79^ (zhanglab.ccmb.med.umich.edu/I-TASSER) was used for 3D protein structure prediction. Predicted structures with a TM-score of over 0.5 were considered reliable as suggested in the I-TASSER user guidelines^80^. TM-align^81^, as implemented in I-TASSER^79^, was used to overlay the predicted protein structure with existing published protein structures.

### Hypnozoite Gene Search

Using the OrthoMCL^68^ clustering between all sequenced *Plasmodium* species used for the phylogenetic analysis (see above), we examined clusters containing only *P. vivax* P01 genes, P. cynomolgi^31^ genes and genes of both of the *P. ovale* species.

### Gene Family Analysis

All *P. malariae, P. ovale*, and *P. vivax* P01 genes were compared pairwise using BLASTP^57^, with genes having a minimum local BLAST hit of 50% identity over 150 amino acids or more being considered connected. These gene connections were visualized in Gephi^82^ using a Fruchterman-Reingold^83^ layout and with unconnected genes.

*P. malariae, P. o. curtisi* and *P. o. wallikeri* protein sequences for *Plasmodium* interspersed repeat *(pir)* genes, excluding pseudogenes, were combined with those from *P. vivax* P01, *P. knowlesi^4^, P. chabaudi* AS v3 (genedb.org/Homepage/Pchabaudi), *P.yoelii 17Xv234*, and *P. berghei v3* (genedb.org/Homepage/Pberghei). Sequences were clustered using tribeMCL^84^ with blast E-value 0.01 and inflation 2. This resulted in 152 subfamilies. We then excluded clusters with one member. The number of genes per species, in each subfamily were plotted in a heatmap using the heatmap.2 function in ggplots in R-3.1.2.

The *pir* genes from two *P. o. curtisi* and two *P. o. wallikeri* assemblies (two high-quality and two draft assemblies) were compared pairwise using BLASTP^57^ with a 99% identity over a minimum of 150 amino acids cutoff. The gene-gene connections were visualized in Gephi^82^ using a Fruchterman-Reingold^83^ layout after removing unconnected genes.

### Mirror Tree Analysis

Using Artemis^66^, 79 *fam-m* and *fam-l* doublets that were confidently predicted as being paired-up were manually selected based on their dispersal throughout the subtelomeres of different chromosomes. The Mirrortree^85^ web server (http://csbg.cnb.csic.es/mtserver/) was used to construct mirror trees for these ^79^ doublets. 35 doublets with recent branching from another doublet were manually selected to enrich for genes under recent selection. To control for chance signals of co-evolution based on their subtelomeric location, the same methodology was repeated by choosing 79 *pir* genes in close proximity of *fam-m* genes as pseudo-doublets and paired up in the Mirrortree^85^ web server.

### Reticulocyte Binding Protein (RBP) Phylogenetic Plot

Full-length RBP genes were manually inspected using ACT^67^ and verified to either be functional or pseudogenized by looking for sequencing reads in other samples that confirm mutations inducing pre-mature stop codons or frameshifts. All functional RBPs were aligned using MUSCLE^49^ and cleaned using GBlocks^69^. PhyML^73^ was used to construct a phylogenetic tree of the different RBPs. Figtree was used to colour the tree (http://tree.bio.ed.ac.uk/software/figtree/).

### SNP Calling

Additional *P. malariae* (PmMY01, PmID01, PmGN01, PmSL01) and *P. o. curtisi* (PocGH01, PocGH02, PocCR01) samples were mapped back against the reference genomes using SMALT (-y 0.8, −i 300) (Supplementary Table 2). As outgroups, *P. malariae-like* (PmlGA01, PmlGA02) and *P. o. wallikeri* (PowCR01, PowCR02) were also mapped against the *P. malariae* and *P. o. curtisi* genomes respectively. The resulting bam files were merged for either of the two genomes, and GATKs^41^ Unified Genotyper was used to call SNPs from the merged bam files (Supplementary Tables 5 and 6). Per GATKs^41^ best practices, SNPs were filtered by quality of depth (QD > 2), depth of coverage (DP > 10), mapping quality (MQ > 20), and strand bias (FS < 60). Additionally, all sites for which we had missing data for any of the samples or where we had heterozygous calls were filtered away. Finally, we filtered away sites that were masked using Dustmasker^75^ to remove repetitive and difficult to map regions.

### Molecular Evolution Analysis

To calculate the genome-wide nucleotide diversity, we extracted all raw SNPs in the genomes excluding the subtelomeres. We then divided the resulting genome size by the number of raw SNPs specific to the core of the genome. This number was averaged for the four *P. malariae* samples and for the two *P. ovale* samples.

The filtered SNPs were used to morph the reference genomes using BCFtools^74^ for each sample, from which sample-specific gene models were obtained. Nucleotide alignments of each gene were then generated. Codons with alignment positions that were masked using Dustmasker^75^ were excluded. For each alignment (*ie*. gene), we calculated HKA^44^, MK^46^, and Ka/Ks^45^ values, see below. Subtelomeric gene families and pseudogenes were excluded from the analysis. The results were analysed and plotted in RStudio (http://www.rstudio.com/).

For the HKA^44^, we counted the proportion of pairwise nucleotide differences intra-specifically *(ie*. within *P. malariae* and within *P. o. curtisi)* and inter-specifically *(ie*. between *P. malariae* and *P. malariae-like*, between *P. o. curtisi* and *P. o. wallikeri)*. The intraspecific comparisons were averaged to get the genes nucleotide diversity pi and these were divided by the average interspecific comparisons, the nucleotide divergence, to get the HKA ratio (HKAr) for each gene.

The MK test^46^ was performed for each gene by obtaining the number of fixed and polymorphic changes, as well as a p-value, as previously described^86^ and then calculating the skew as log2(((N_poly_+1)/(S_poly_+1))/((N_fix_+1)/(S_fix_+1))) where N_poly_ and N_fix_ are polymorphic and fixed non-synonymous substitutions respectively, while Spoly and Sfix refer to the synonymous substitutions.

To calculate the average Ka/Ks ratio^45^, we took the cleaned alignments of the MK test, extracting the pairwise sequences of *P. malariae* and *P. malariae-like* (and of *P. o. curtisi* and *P. o. wallikeri)*. The Bio::Align::DNAStatistics module was used to calculate the Ka/Ks values for each pair^87^, averaging across samples within a species.

Using existing RNA-Seq data from seven different life-cycle stages in *P. falciparum^25^*, reads were mapped against spliced gene sequences (exons, but not UTRs) from the *P. falciparum* 3D7 reference genome^2^ using Bowtie288 v2.1.0 (-a-X 800-x). Read counts per transcript were estimated using eXpress v1.3.0^89^. Genes with an effective length cutoff below 10 in any sample were removed. Summing over transcripts generated read counts per gene. Each gene in *P. malariae* and *P. ovale* was classified by their *P. falciparum* orthologs maximum expression stage.

### RBP1a Receptor Search

To find the putative RBP1a receptor, we performed an OrthoMCL^68^ clustering between Human, Chimpanzee^90^, and common marmoset^91^ genes. The common marmoset has been found infected with *P. brasilianum (P. malariae)* in the wild^92^. Genes without transmembrane domain as well as those annotated as ‘predicted’ were removed. To remove false positive, all remaining genes were searched against the Chimpanzee genes using BLASTP^57^ with a threshold of 1e-10.

## Acknowledgements

This work was supported by the Medical Research Council [MR/J004111/1] [MR/L008661/1] and the Wellcome Trust [098051]. S.A. and R.N.P. are funded by the Wellcome Trust (Senior Fellowship in Clinical Science awarded to RNP, 091625). F.R., B.O., and F.P. are financed by the ANR JCJC 2012 ORIGIN, the LMI Zofac, as well as by CNRS, IRD, and CIRMF. C.I.N. is funded by the Wellcome Trust [104792]. A.A.D. is funded as a Sanger International Fellow. The authors thank Eric Willaume from the Park of La Lékédi, and the different people involved in the sanitary controls of the chimpanzees. J.S.M. and N.M.A. are supported by NHMRC Practitioner Fellowships (#104072).

## Author Contributions

G.G.R. carried out the sequence assembly, genome annotation and all the data analysis; U.C.B. performed manual gene curation; M.S. coordinated sequencing; A.J.R., M. M., and F.P. performed data analysis; G.G.R., T.O.A., L.AE., J.W.B., D.P.K., C.I.N., M.B., and T.D.O. designed the *P. ovale* project; G.G.R., F.R., B.O., F.P., C.I.N., M.B., and T.D.O. designed the *P. malariae-like* project; G.G.R., A.A.D., O.M.A, N.M.A., S.A., R.N.P., J.S.M., C.I.N., M.B., and T.D.O. designed the *P. malariae* project; G.G.R., C.I.N., M.B., T.D.O wrote the manuscript; All authors read and critically revised the manuscript; and C.I.N., M.B., T.D.O. directed the overall study.

**Supplementary Figure 1.**
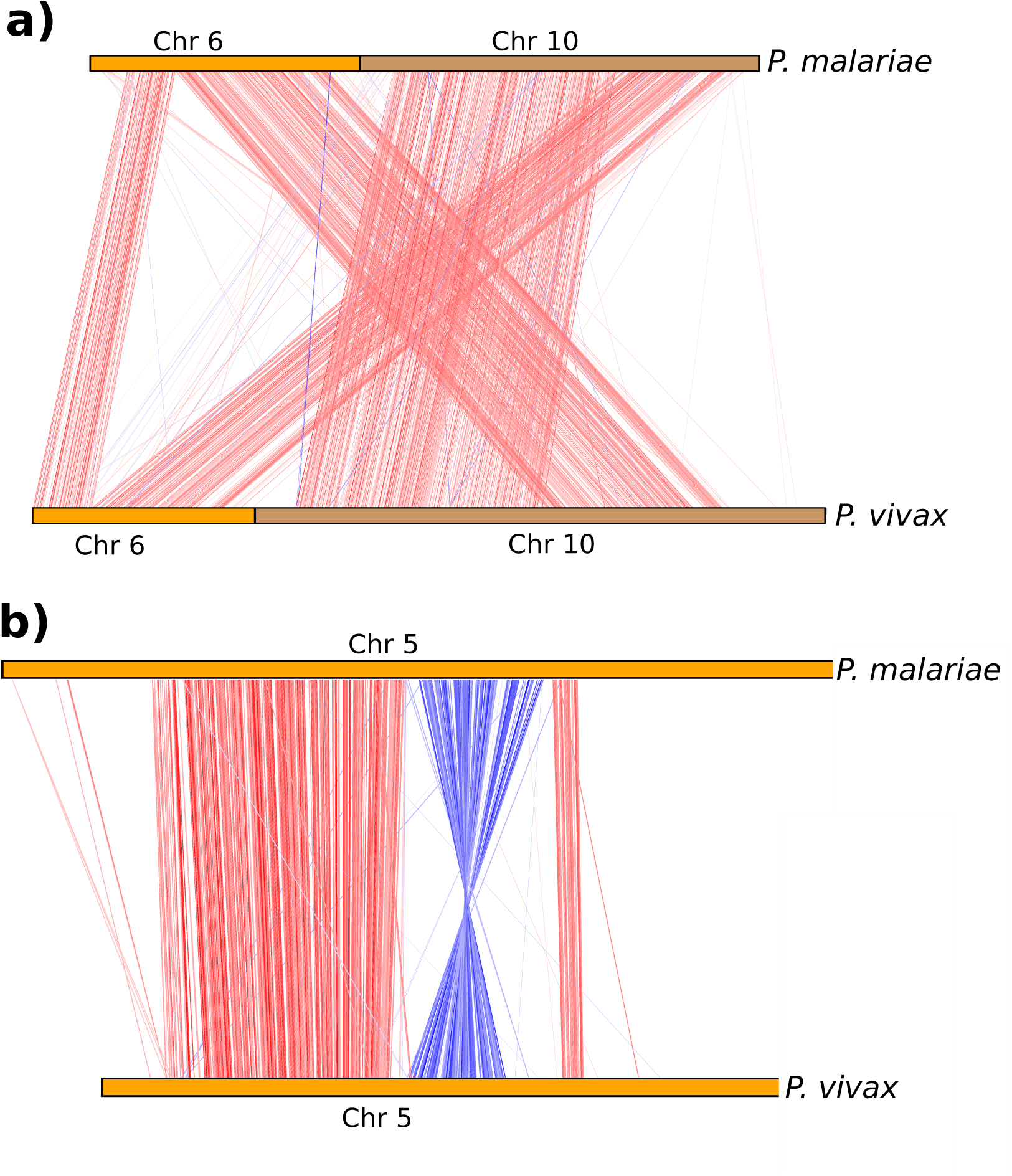
Recombination breakpoints in *P. malariae* compared to *P. vivax*. **a)** ACT^67^ view showing recombination of chromoso 0 in *P malariae*. The red lines indicate blast similarities, chromosome e and chromosome 10 in brown. **b)** ACT^67^ view showing internal inversi some 5 of *P. malariae*. Red lines indicate blast similarities and blue line erted blast hits.

**Supplementary Figure 2.**
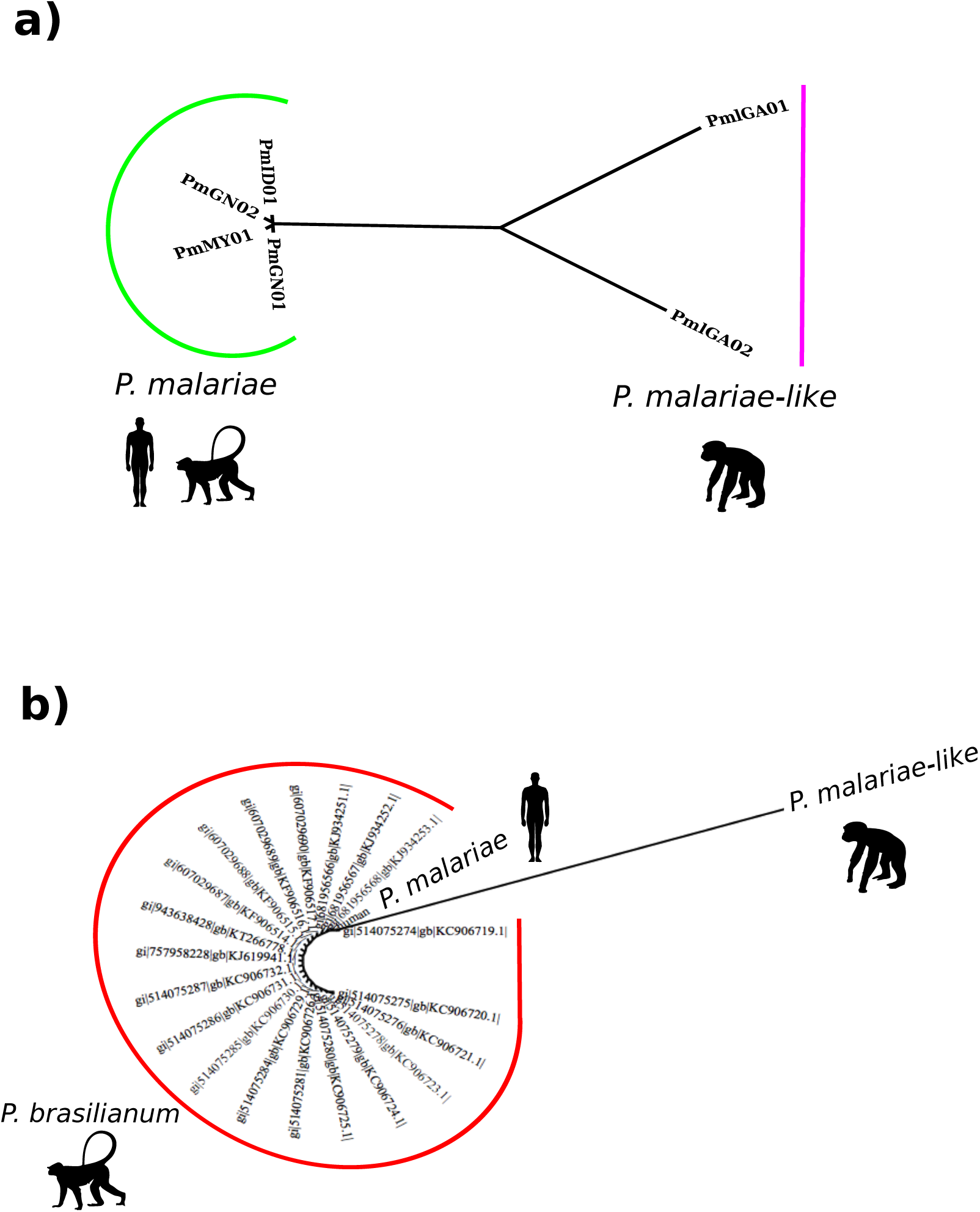
*P. malariae-like* has significantly longer branch lengths than *P. malariae*, and *P. brasilianum* is identical to *P. malariae*. **a)** A phylogenetic tree of all *P. malariae* and *P. malariae-like* samples generated using PhyML73 based on all *P. malariae* genes. *P. malariae* samples are indicated by a green bar and *P. malariae-like* samples are indicated by a purple bar. Silhouettes represent infectivity. **b)** A PhyML_ENREF_72^73^ phylogenetic tree of all *P. brasilianum* 18S rRNA sequences^21^, indicated by a red bar, and the corresponding 18S rRNA sequences from the *P. malariae* and *P. malariae-like* assemblies, labeled as such. Silhouettes represent the host origin for each sample.

**Supplementary Figure 3.**
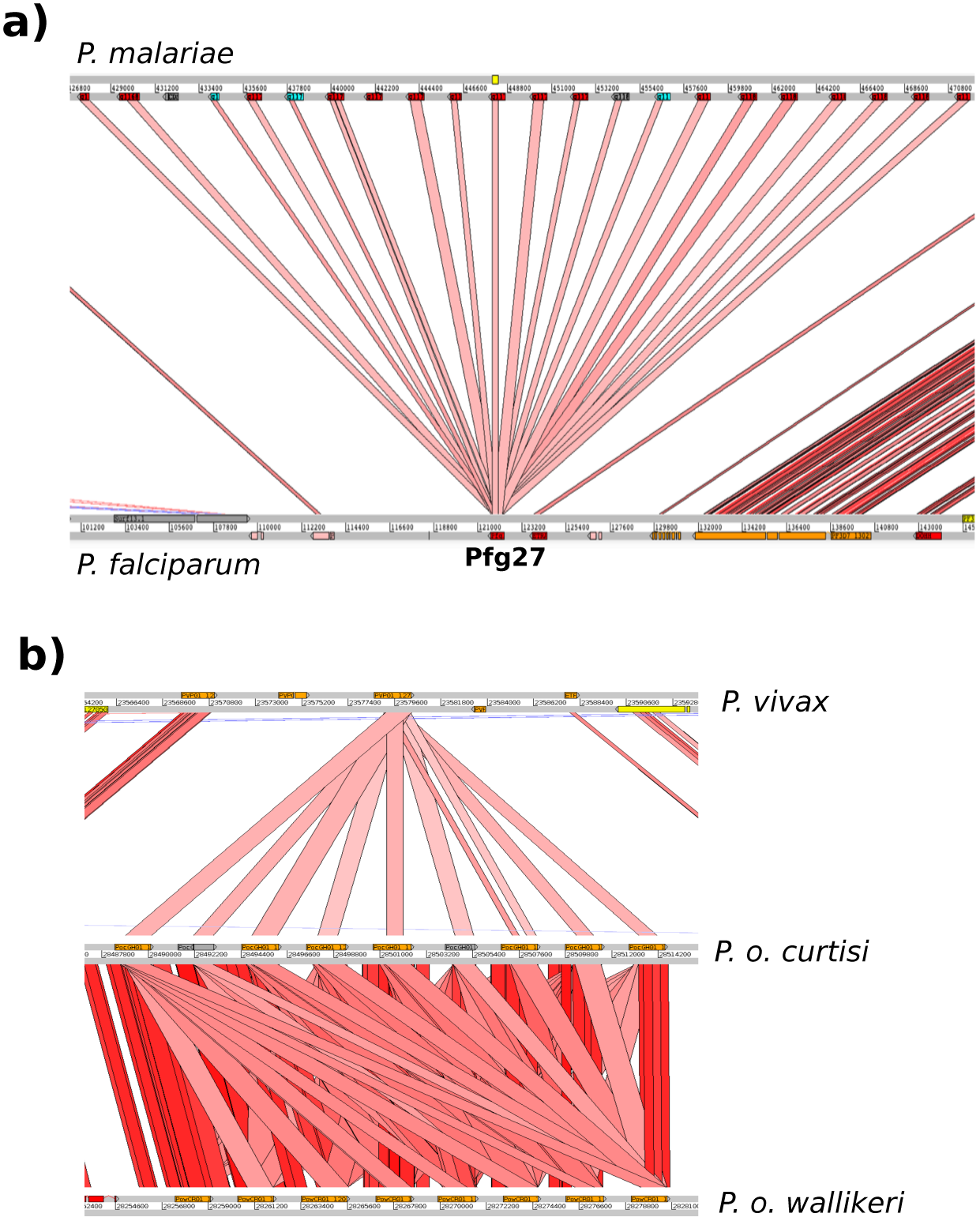
Large gene duplications in *P. malariae* and *P. ovale*. **a)** Expansion of Pfg27 in *P. malariae* (top) compared to *P. falciparum* (bottom) with red lines indicating blast similarities. Functional genes are in red and pseudogenes in grey. **b)** Expansion of PVP01_1270800 (PF3D7_1475900 in *P falciparum)*, a gene with no known function, in *P. o. curtisi* and *P. o. wallikeri*, with different copy numbers in each, compared to the one copy in *P. vivax*. Functional genes shown in orange and pseudogenes shown in grey.

**Supplementary Figure 4.**
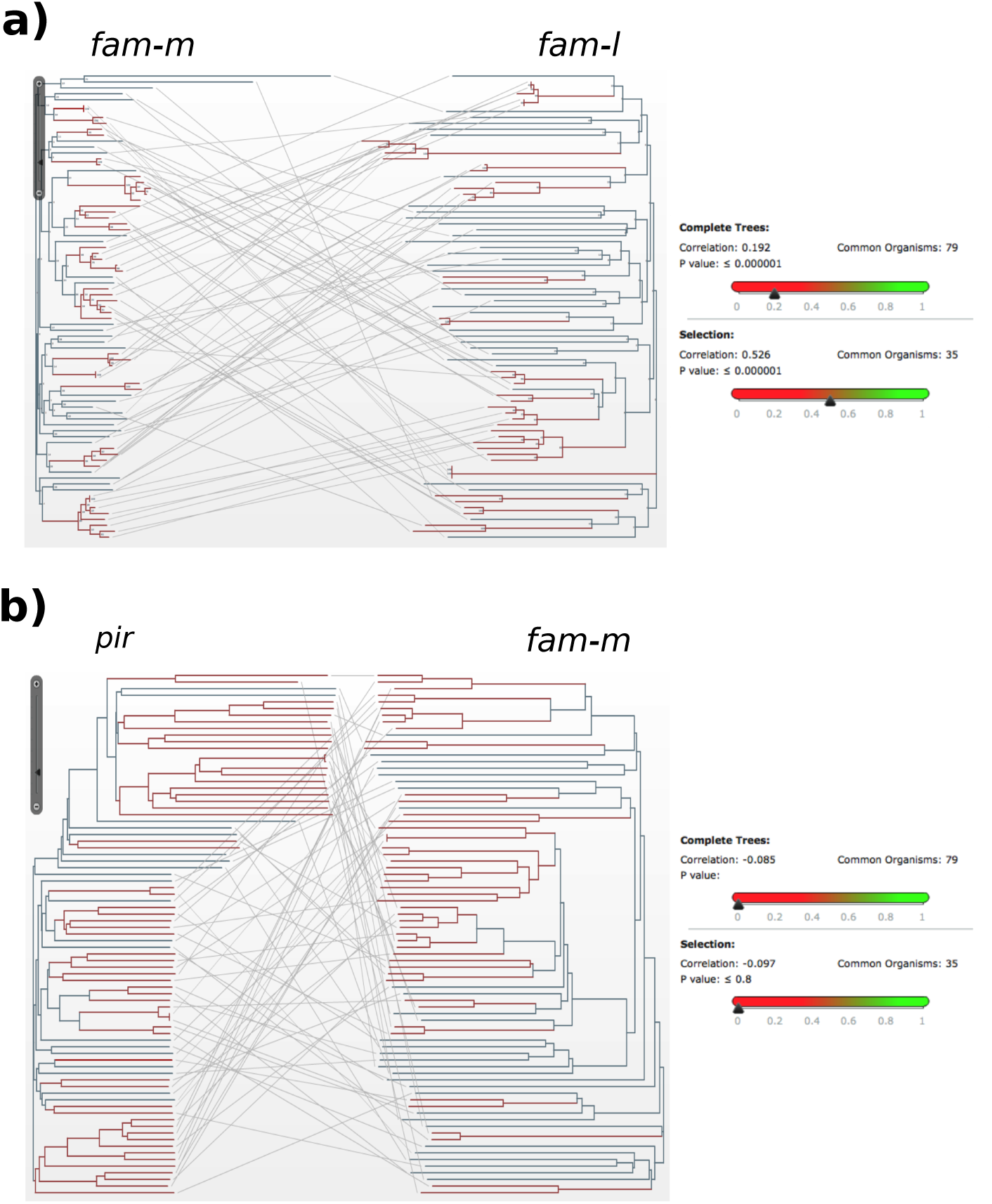
Co-evolution of *fam-m* and *fam-l* genes, but not with *pir* genes. **a)** Mirror tree^85^ for 79 *fam-m* and *fam-l* doublets, where the two phylogenetic trees correspond to either of the families with lines connecting branch tips of the same doublet. 35 branches (Red) were manually selected due to exhibiting recent branching. Inset shows the correlation between the two trees for all branches (above, r^2^=0.19, p < 0.001) and red branches (below, r^2^=0.53, p < 0.001). This shows that the two families are co-evolving, especially when doublets that recently branched are selected. **b)** Mirror tree^85^ for 79 *pir* and *fam-m* pseudo-doublets (Methods), where the two phylogenetic trees correspond to either of the families with lines connecting branch tips of the same doublet. 35 branches (Red) were manually selected due to exhibiting recent branching. Inset shows the correlation between the two trees for all branches (above, r^2^=0.09, p > 0.05) and red branches (below, r^2^=0.10, p > 0.05). This shows that the two families are not co-evolving, and that subtelomeric location does not produce sporadic signals of co-evolution.

**Supplementary Figure 5.**
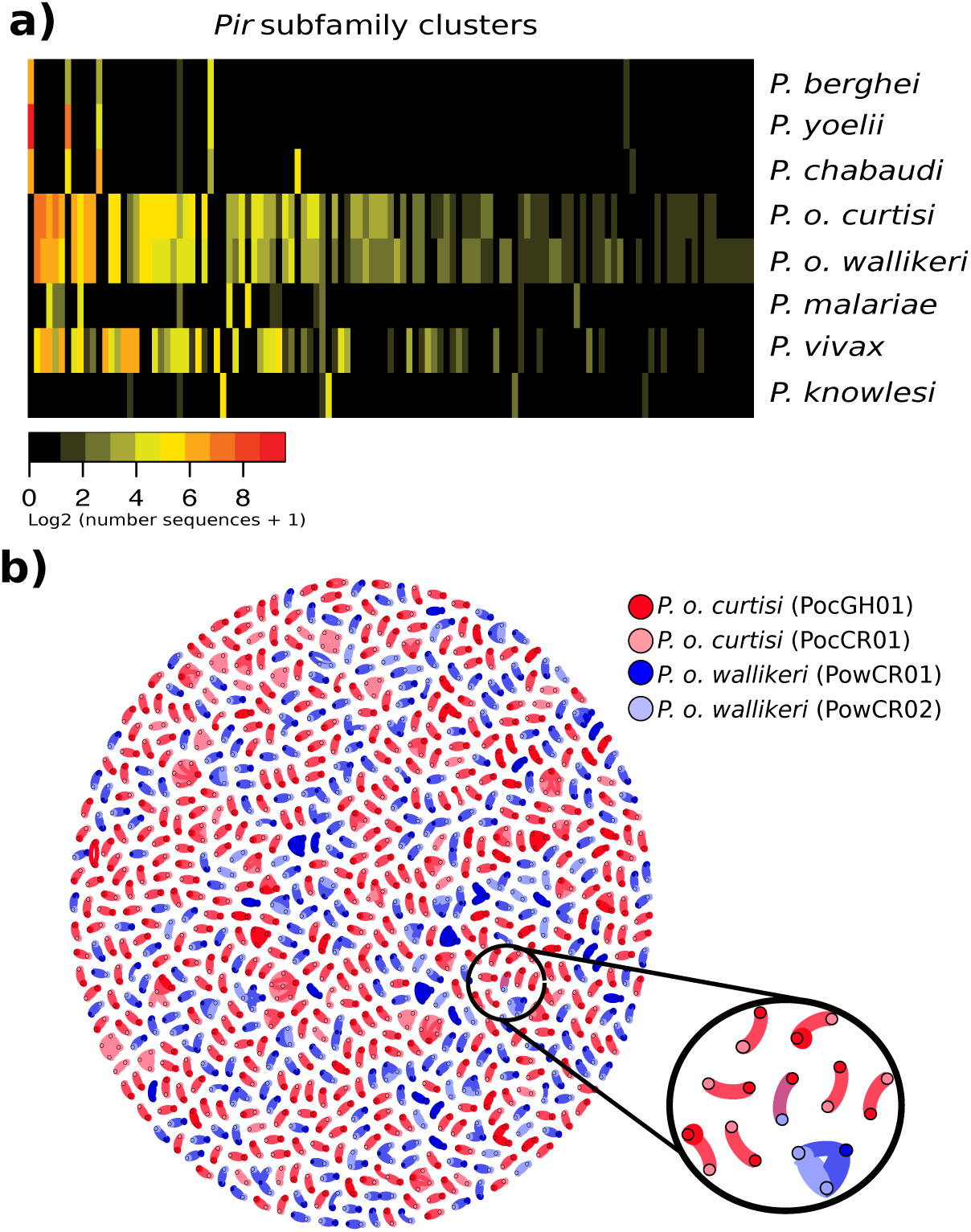
*Pir* genes in *P. malariae* and *P. ovale* resemble those in *P. vivax*, and *pir* gense are less similar between the two *P. ovale* than within. **a)** Heatmap showing the sharing of *pir* subfamilies between different species based on tribeMCL^84^. Columns show *pir* subfamilies and rows show species. Colours indicate the number of genes classified into each subfamily for each species. Subfamilies were ordered by size, species were ordered for clarity. *Pir* genes in rodent-infecting species fall into a small number of well-defined families. Those in *P. vivax, P. malariae* and *P. ovale* are however much more diverse. There is little overlap between rodent subfamilies and human-infecting subfamilies, despite *P. ovale* being a sister taxa to the rodent-infecting species. *P. knowlesi* has some sharing with other species, but its largest families are species-specific, suggesting it has undergone specialization of its *pir* repertoire. **b)** Gene network of *pir* genes for both high-quality assemblies of *P. o. curtisi* (Dark red) and *P. o. wallikeri* (Dark blue) and draft assemblies of each (Light red and light blue respectively). *Pir* genes with BLASTP57 identity hits of 99%+ over 150 amino acids become connected in the graph. Genes without connections were removed. There is 1 connection between the two species (circled in black and with a zoomed in version), 801 between the *P. o. curtisi* assemblies, 524 between the *P. o. wallikeri* assemblies, 527 on average within each *P. o. curtisi* assembly, and 423 on average within each *P. o. wallikeri* assembly.

**Supplementary Figure 6.**
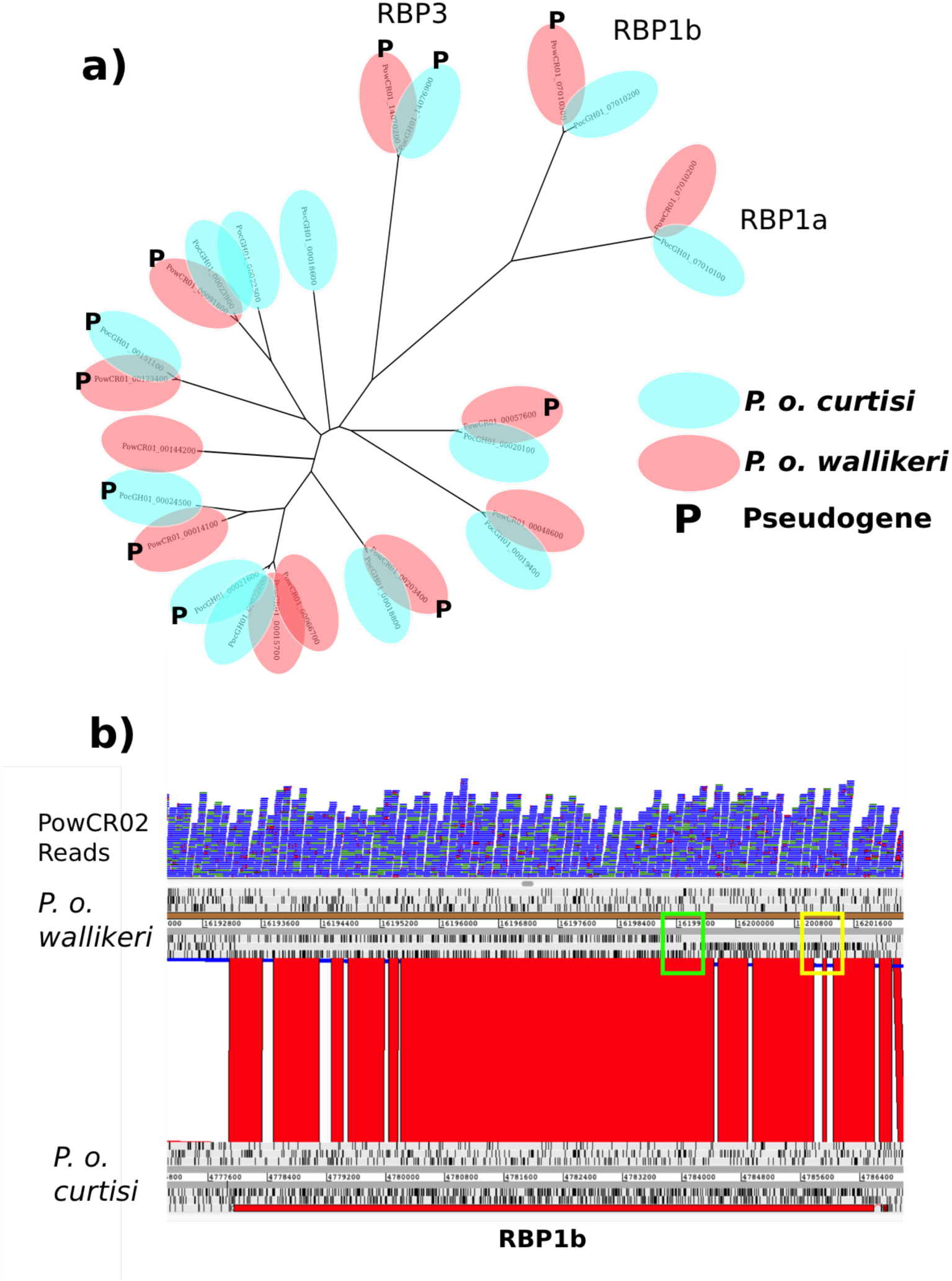
Multiple RBP genes are pseudogenized between the two *P. ovale* species. **a)** PhyML^73^ generated phylogenetic tree of all RBP genes over 1kb long in *P. o. curtisi* (light blue) and *P. o. wallikeri* (light red). Pseudogenes are denoted with **P**. Multiple functional RBP2 genes match up with pseudogenized copies in the other genome. **b)** ACT67 view of RBP1b in red for *P. o. curtisi* (bottom) and the corresponding disrupted open reading frame in *P. o. wallikeri* (top), with black ticks indicating stop codons. Reads (in blue) from an additional *P. o. wallikeri* sample (PowCR02) confirm the bases introducing the frameshift (green square) and premature stop codon (yellow square) in RBP1b.

**Supplementary Table 1.** **Showing the number of samples positive for different *Plasmodium* species in the Pf3K dataset**.

| Country | Total Samples | *P. falciparum* Positive | *P. vivax* Positive | *P. malariae* Positive | *P. ovale* Positive | *P. knowlesi* Positive |
| --- | --- | --- | --- | --- | --- | --- |
| The Gambia | 65 | 65 | 0 | 0 | 0 | 0 |
| Guinea | 100 | 100 | 0 | 7 | 3 | 0 |
| Thailand | 148 | 148 | 11 | 0 | 0 | 0 |
| Ghana | 617 | 617 | 5 | 12 | 9 | 0 |
| Cambodia | 570 | 570 | 50 | 0 | 0 | 0 |
| Mali | 96 | 96 | 0 | 1 | 0 | 0 |
| Bangladesh | 50 | 50 | 4 | 0 | 0 | 0 |
| Malawi | 369 | 369 | 4 | 4 | 4 | 0 |
| Vietnam | 97 | 97 | 16 | 0 | 0 | 0 |
| Myanmar | 60 | 60 | 7 | 0 | 0 | 0 |
| Laos | 85 | 85 | 4 | 0 | 2 | 0 |
| DR Congo | 113 | 113 | 1 | 2 | 1 | 0 |
| Nigeria | 5 | 5 | 0 | 0 | 0 | 0 |
| Senegal | 137 | 137 | 0 | 0 | 1 | 0 |
| **GLOBAL** | 2512 | 2512 | 102 | 26 | 19 | 0 |

The first column shows country of origin for the different samples, with the second column showing the total number of samples collected in that country. The following five columns show the number of these samples that are positive for the different *Plasmodium* species. All samples are positive for *P. falciparum*, which is expected because all the samples were initially identified as *P. falciparum*. We do not see any samples positive for *P. knowlesi*, because it has a very limited geographic range and isnt found in any of the sampled countries.

**Supplementary Table 2.** **Showing the sample origins, sequencing statistics, and uses in this study**

| Species | Sample ID | Accession Number | Origin | Sequencing Platform | Read Length | Usage | Library type |
| --- | --- | --- | --- | --- | --- | --- | --- |
| *P. malariae* | PmUG01 | ERS1110315 | Uganda | PacBio RS II P6/C4 | N/A | Reference Genome | Size-selected |
|  | PmMY01 | ERS1110317 | Malaysia | Illumina MiSeq v2<br>Illumina HiSeq 2000 v4 | 150bp PE<br>125bp PE | SNP Calling | Amplification free |
|  | PmID01 | ERS1110321 | Papua Indonesia | Illumina MiSeq v2<br>Illumina HiSeq 2000 v4 | 150bp PE<br>125bp PE | SNP Calling | Amplification free |
|  | PmGN01 | ERS1110325 | Mali | Illumina MiSeq v2 | 150bp PE | SNP Calling | Standard |
|  | PmGN02 | ERS567899 | Guinea | Illumina MiSeq v2 | 150bp PE<br>250bp PE | SNP Calling | Standard |
| *P. malariae*-like | PmlGA01 | ERS434571 | Gabon | Illumina MiSeq v2 | 150bp PE | Draft Genome | WGA - Amplification free |
|  | PmlGA02 | ERS434565 | Gabon | Illumina MiSeq v2 | 150bp PE | SNP Calling | WGA - Amplification free |
| *P. ovale curtisi* | PocGH01 | ERS013096 | Ghana | Illumina HiSeq 2000 v3 | 100bp PE | Reference Genome | Standard |
|  | PocGH02 | ERS360497 | Ghana | Illumina HiSeq 2000 v3 | 100bp PE | SNP Calling | Standard |
|  | PocCR01 | ERS418861 | Cameroon | Illumina HiSeq 2000 v3 | 100bp PE | SNP Calling | Standard |
| *P. ovale wallikeri* | PowCR01 | ERS418894 | Cameroon | Illumina HiSeq 2000 v3 | 100bp PE | Draft Genome | Standard |
|  | PowCR02 | ERS418932 | Cameroon | Illumina HiSeq 2000 v3 | 100bp PE | SNP Calling | Standard |

**Supplementary Table 3.**
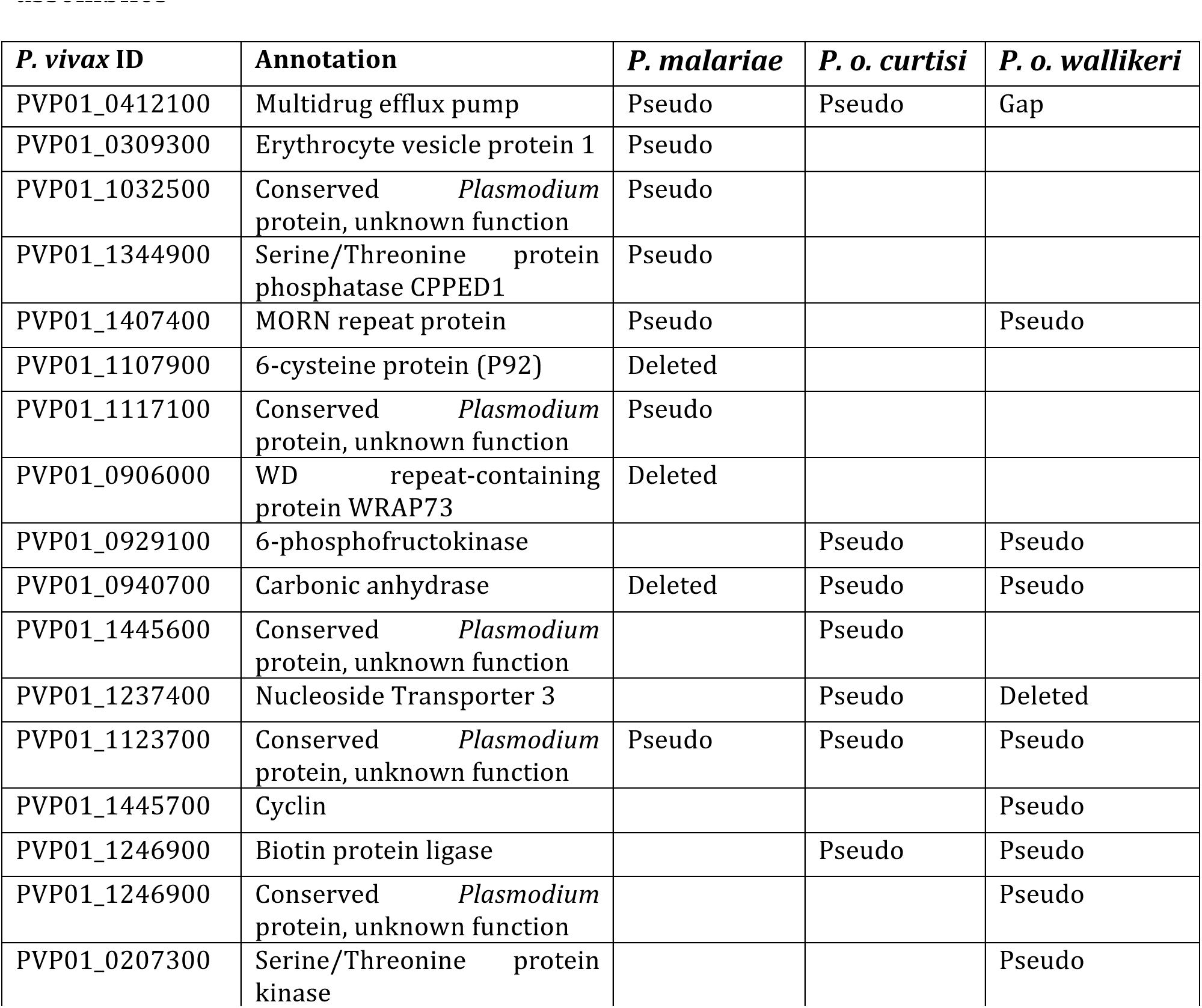
**Showing pseudogenized and deleted core genes in the three high-quality assemblies**

The first column shows the gene ID of the *P. vivax* P01 homolog of the gene pseudogenized/deleted in one or more of the three human malaria parasite assemblies. The second column is the *P. vivax* P01 annotation of that gene. The following three columns show whether the gene is functional (blank), pseudogenized (Pseudo), deleted (Deleted), or missing due to a sequencing gap (Gap).

**Supplementary Table 4.** **Showing potential hypnozoite genes**

| PVP01 Annotation | *P. vivax* | *P. cynomolgi* | *P. o. curtisi* | *P. o. wallikeri* |
| --- | --- | --- | --- | --- |
| Ring-exported protein 4* | PVP01_0623900 | Pcyb_063280 | PocGH01_00129400 | PowCR01_06029200 |
| Conserved Plasmodium protein | PVP01_1402600 | Pcyb_141110 | PocGH01_00080600 | PowCR01_00100500 |

\*not in the same orthologous group as *P. falciparum* REX4

These are the two orthoMCL gene clusters that contain exclusively all hypnozoite-forming *Plasmodium* species and are not part of subtelomeric gene families.

**Supplementary Table 5.** ***P. malariae* SNP Calling Results**

| Species | *P. malariae* |  |  |  | *P. malariae-like* |  |
| --- | --- | --- | --- | --- | --- | --- |
| Sample ID | PmMY01 | PmID01 | PmGN01 | PmGN02 | PmlGA01 | PmlGA02 |
| **Raw SNPs** | 218334 | 164541 | 173028 | 239655 | 458790 | 211686 |
| **- Private** | 48094 | 19475 | 25901 | 66377 | 260540 | 68793 |
| **- Ref** | 712758 | 696634 | 706817 | 737236 | 386915 | 261042 |
| **- Missing\*** | 50394 | 86900 | 74936 | 28776 | 165813 | 415250 |
| **Filtered SNPs** | 8970 | 8589 | 7742 | 7878 | 161551 | 140113 |
| **- Private** | 2149 | 2066 | 1908 | 2066 | 77781 | 56571 |
| **- Ref** | 221923 | 222247 | 223058 | 223003 | 69466 | 90571 |
| **- Missing\*** | 0 | 0 | 0 | 0 | 0 | 0 |

\*sites at which the sample has no coverage

SNP calling results as per mapping all *P. malariae* and *P. malariae-like* samples against the PmUG01 PacBio reference genome assembly. The raw SNPs are the total number of SNPs that we call using GATK default parameters in the different samples. Of these raw SNPs, some are exclusive to a certain sample (Private), are identical to the reference genome (Ref), or there is no coverage and therefore no SNP call could be made (Missing). The same information is also shown for the filtered SNPs, which were filtered according to a number of different parameters (Methods).

**Supplementary Table 6.** ***P. ovale* SNP Calling Results**

| **Species** | ***P. o. curtisi*** |  | ***P. o. wallikeri*** |  |
| --- | --- | --- | --- | --- |
| **Sample ID** | **PocGH02** | **PocCR01** | **PowCR01** | **PowCR02** |
| **Raw SNPs** | 171465 | 277978 | 2139946 | 1881088 |
| **- Private** | 36487 | 99083 | 333727 | 83609 |
| **- Ref** | 2287008 | 2249682 | 693405 | 674071 |
| **- Missing\*** | 84743 | 72495 | 104013 | 149166 |
| **Filtered SNPs** | 29099 | 46695 | 1415164 | 1410434 |
| **- Private** | 6162 | 16026 | 21081 | 16699 |
| **- Ref** | 1433387 | 1416042 | 45978 | 50845 |
| **- Missing\*** | 0 | 0 | 0 | 0 |

\*sites at which the sample has no coverage

SNP calling results as per mapping all *P. o. curtisi* and *P. o. wallikeri* samples against the PocGH01 Illumina reference genome assembly. The raw SNPs are the total number of SNPs that we call using GATK default parameters in the different samples. Of these raw SNPs, some are exclusive to a certain sample (Private), are identical to the reference genome (Ref), or there is no coverage and therefore no SNP call could be made (Missing). The same information is also shown for the filtered SNPs, which were filtered according to a number of different parameters (Methods).

**Supplementary Table 7.** **Genes with significant scores in two or more population genetics measures**

| Species | Gene ID | Gene Product |
| --- | --- | --- |
| *P. malariae* | PmUG01_10020100 | Heat shock protein 90, putative |
| *P. malariae* | PmUG01_10046100 | Methione-tRNA ligase, putative |
| *P. malariae* | PmUG01_07021900 | Conserved Plasmodium protein, unknown function |
| *P. malariae* | PmUG01_09048000 | Beta-catenin-like protein 1, putative |
| *P. malariae* | PmUG01_13021900 | P-type ATPase4, putative |
| *P. malariae* | PmUG01_07035300 | Thioredoxin reductase, putative |
| *P. malariae* | PmUG01_10046000 | Conserved Plasmodium protein, unknown function |
| *P. malariae* | PmUG01_11017800 | Conserved Plasmodium protein, unknown function |
| *P. malariae* | PmUG01_04014000 | DEAD/DEAH box ATP-dependent RNA helicase, putative |
| *P. malariae* | PmUG01_07042000 | Merozoite surface protein 1, putative |
| *P. malariae* | PmUG01_10013600 | Formin-1, putative |
| *P. malariae* | PmUG01_13030700 | rRNA (adenosine-2'-O)-methyltransferase |
| *P. malariae* | PmUG01_12012900 | Conserved Plasmodium protein, unknown function |
| *P. o. curtisi* | PocGH01_00022600 | Reticulocyte binding protein 2b, putative |

For the three population genetics measures (HKAr, Ka/Ks, and Skew), the table shows that genes that have significant value in two or more of these measures. These genes therefore represent genes under significant selection pressures.

**Supplementary Table 8.**
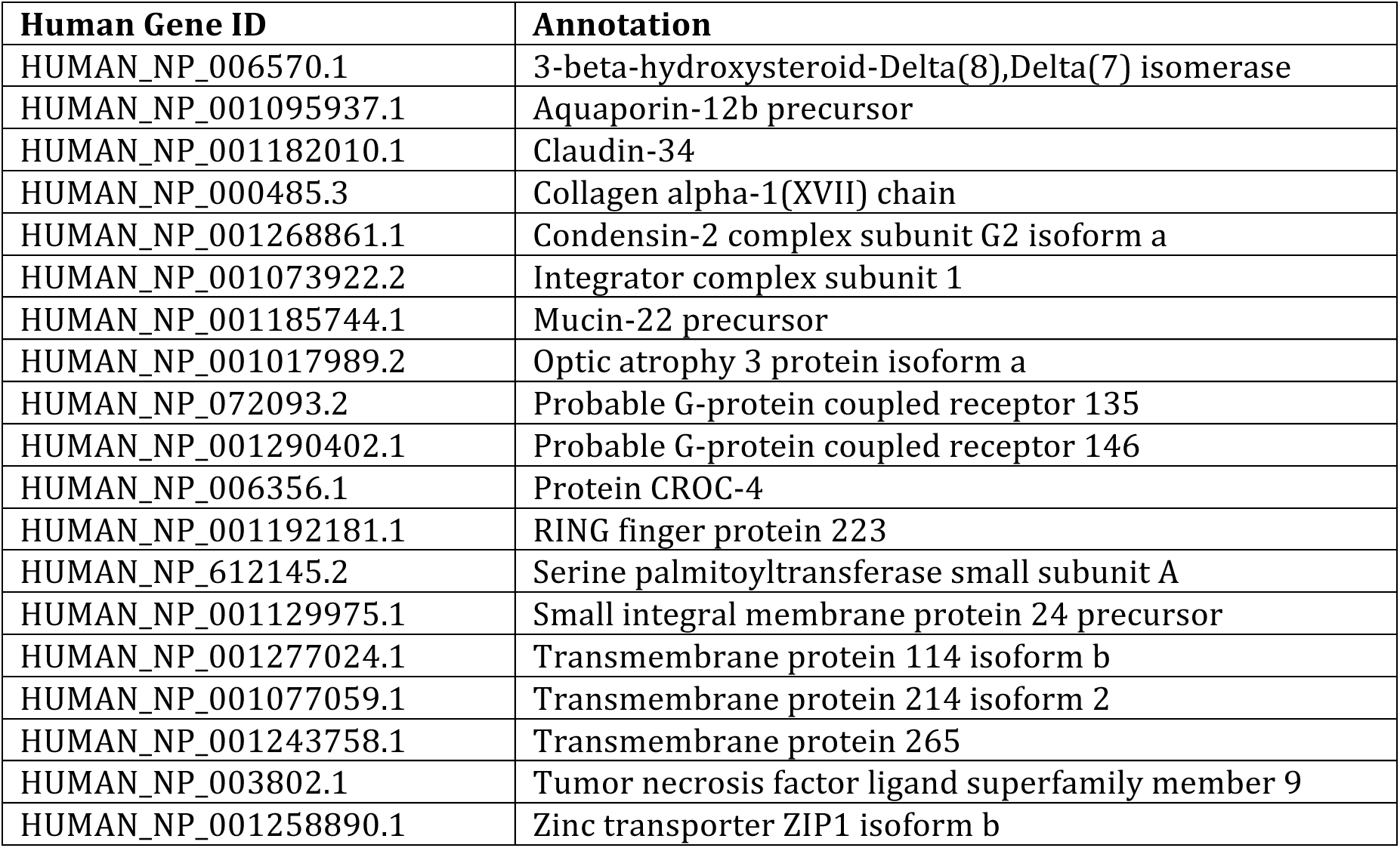
**List of Potential RBP1a receptors**

| Human Gene ID | Annotation |
| --- | --- |
| HUMAN_NP_006570.1 | 3-beta-hydroxysteroid-Delta(8),Delta(7) isomerase |
| HUMAN_NP_001095937.1 | Aquaporin-12b precursor |
| HUMAN_NP_001182010.1 | Claudin-34 |
| HUMAN_NP_000485.3 | Collagen alpha-1(XVII) chain |
| HUMAN_NP_001268861.1 | Condensin-2 complex subunit G2 isoform a |
| HUMAN_NP_001073922.2 | Integrator complex subunit 1 |
| HUMAN_NP_001185744.1 | Mucin-22 precursor |
| HUMAN_NP_001017989.2 | Optic atrophy 3 protein isoform a |
| HUMAN_NP_072093.2 | Probable G-protein coupled receptor 135 |
| HUMAN_NP_001290402.1 | Probable G-protein coupled receptor 146 |
| HUMAN_NP_006356.1 | Protein CROC-4 |
| HUMAN_NP_001192181.1 | RING finger protein 223 |
| HUMAN_NP_612145.2 | Serine palmitoyltransferase small subunit A |
| HUMAN_NP_001129975.1 | Small integral membrane protein 24 precursor |
| HUMAN_NP_001277024.1 | Transmembrane protein 114 isoform b |
| HUMAN_NP_001077059.1 | Transmembrane protein 214 isoform 2 |
| HUMAN_NP_001243758.1 | Transmembrane protein 265 |
| HUMAN_NP_003802.1 | Tumor necrosis factor ligand superfamily member 9 |
| HUMAN_NP_001258890.1 | Zinc transporter ZIP1 isoform b |

The first column shows the 19 transmembrane-containing human genes that are shared between humans and the common marmoset, but not with chimpanzees. As RBP1a is the only RBP with large differences between *P. malariae* and *P. malariae-like*, these genes may represent interesting RBP1a receptor targets.

## References

1 Keeling, P. J. & Rayner, J. C. The origins of malaria: there are more things in heaven and earth. Parasitology 142 Suppl 1, S16–25, doi:10.1017/S0031182014000766(2015).

2 Gardner, M. J. et al. Genome sequence of the human malaria parasite Plasmodium falciparum. Nature 419, 498–511, doi:10.1038/nature01097 (2002).

3 Carlton, J. M. et al. Comparative genomics of the neglected human malaria parasite Plasmodium vivax. Nature 455, 757–763, doi:10.1038/nature07327 (2008).

4 Pain, A. et al. The genome of the simian and human malaria parasite Plasmodium knowlesi. Nature 455, 799–803, doi:10.1038/nature07306 (2008).

5 Chin, W., Contacos, P. G., Coatney, G. R. & Kimball, H. R. A Naturally Acquited Quotidian-Type Malaria in Man Transferable to Monkeys. Science 149, 865 (1965).

6 Cunningham, D., Lawton, J., Jarra, W., Preiser, P. & Langhorne, J. The pir multigene family of Plasmodium: antigenic variation and beyond. Mol Biochem Parasitol 170, 65–73, doi:10.1016/j.molbiopara.2009.12.010 (2010).

7 Cowman, A. F. & Crabb, B. S. The Plasmodium falciparum genome--a blueprint for erythrocyte invasion. Science 298, 126–128, doi:10.1126/science.1078169(2002).

8 Mu, J. et al. Genome-wide variation and identification of vaccine targets in the Plasmodium falciparum genome. Nat Genet 39, 126–130, doi:10.1038/ng1924 (2007).

9 Roucher, C., Rogier, C., Sokhna, C., Tall, A. & Trape, J. F. A 20-year longitudinal study of Plasmodium ovale and Plasmodium malariae prevalence and morbidity in a West African population. PLoS One 9, e87169, doi:10.1371/journal.pone.0087169(2014).

10 Doderer-Lang, C. et al. The ears of the African elephant: unexpected high seroprevalence of Plasmodium ovale and Plasmodium malariae in healthy populations in Western Africa. Malar J 13, 240, doi:10.1186/1475-2875-13-240 (2014).

11 Bousema, T., Okell, L., Felger, I. & Drakeley, C. Asymptomatic malaria infections: detectability, transmissibility and public health relevance. Nat Rev Microbiol 12, 833–840, doi:10.1038/nrmicro3364 (2014).

12 Collins, W. E. & Jeffery, G. M. Plasmodium ovale: parasite and disease. Clin Microbiol Rev 18, 570–581, doi:10.1128/CMR.18.3.570-581.2005 (2005).

13 Collins, W. E. & Jeffery, G. M. Plasmodium malariae: parasite and disease. Clin Microbiol Rev 20, 579–592, doi:10.1128/CMR.00027-07 (2007).

14 Langford, S. et al. Plasmodium malariae Infection Associated with a High Burden of Anemia: A Hospital-Based Surveillance Study. PLoS Negl Trop Dis 9, e0004195, doi:10.1371/journal.pntd.0004195 (2015).

15 Siala, E. et al. [Relapse of Plasmodium malariae malaria 20 years after living in an endemic area]. Presse Med 34, 371–372 (2005).

16 Sutherland, C. J. et al. Two nonrecombining sympatric forms of the human malaria parasite Plasmodium ovale occur globally. J Infect Dis 201, 1544–1550, doi:10.1086/652240 (2010).

17 Arisue, N. et al. The Plasmodium apicoplast genome: conserved structure and close relationship of P. ovale to rodent malaria parasites. Mol Biol Evol 29, 2095–2099, doi:10.1093/molbev/mss082 (2012).

18 Sundararaman, S. A. et al. Genomes of cryptic chimpanzee Plasmodium species reveal key evolutionary events leading to human malaria. Nat Commun 7, 11078, doi:10.1038/ncomms11078 (2016).

19 Gronau, I., Hubisz, M. J., Gulko, B., Danko, C. G. & Siepel, A. Bayesian inference of ancient human demography from individual genome sequences. Nat Genet 43,1031–1034, doi:10.1038/ng.937 (2011).

20 Silva, J. C., Egan, A., Arze, C., Spouge, J. L. & Harris, D. G. A new method for estimating species age supports the coexistence of malaria parasites and their Mammalian hosts. Mol Biol Evol 32, 1354–1364, doi:10.1093/molbev/msv005(2015).

21 Lalremruata, A. et al. Natural infection of Plasmodium brasilianum in humans: Man and monkey share quartan malaria parasites in the Venezuelan Amazon. EBioMedicine 2, 1186–1192, doi:10.1016/j.ebiom.2015.07.033 (2015).

22 Guimaraes, L. O. et al. Merozoite surface protein-1 genetic diversity in Plasmodium malariae and Plasmodium brasilianum from Brazil. BMC Infect Dis 15, 529, doi:10.1186/s12879-015-1238-8 (2015).

23 Lobo, C. A., Fujioka, H., Aikawa, M. & Kumar, N. Disruption of the Pfg27 locus by homologous recombination leads to loss of the sexual phenotype in P. falciparum. Mol Cell 3, 793–798 (1999).

24 Olivieri, A. et al. The Plasmodium falciparum protein Pfg27 is dispensable for gametocyte and gamete production, but contributes to cell integrity during gametocytogenesis. Mol Microbiol 73, 180–193, doi:10.1111/j.1365-2958.2009.06762.x (2009).

25 Lopez-Barragan, M. J. et al. Directional gene expression and antisense transcripts in sexual and asexual stages of Plasmodium falciparum. BMC Genomics 12, 587, doi:10.1186/1471-2164-12-587 (2011).

26 Zhang, Y. Template-based modeling and free modeling by I-TASSER in CASP7. Proteins 69 Suppl 8, 108–117, doi:10.1002/prot.21702 (2007).

27 Wu, R. Control of glycolysis by phosphofructokinase in slices of rat liver, Novikoff hepatoma, and adenocarcinomas. Biochem Biophys Res Commun 14, 79–85 (1964).

28 Nolder, D. et al. An observational study of malaria in British travellers: Plasmodium ovale wallikeri and Plasmodium ovale curtisi differ significantly in the duration of latency. BMJ Open 3, doi:10.1136/bmjopen-2013-002711 (2013).

29 Roques, M. et al. Plasmodium P-Type Cyclin CYC3 Modulates Endomitotic Growth during Oocyst Development in Mosquitoes. PLoS Pathog 11, e1005273, doi:10.1371/journal.ppat.1005273 (2015).

30 Claudio, J. O. et al. Cloning and expression analysis of a novel WD repeat gene, WDR3, mapping to 1p12-p13. Genomics 59, 85–89, doi:10.1006/geno.1999.5858(1999).

31 Tachibana, S. et al. Plasmodium cynomolgi genome sequences provide insight into Plasmodium vivax and the monkey malaria clade. Nat Genet 44, 1051–1055, doi:10.1038/ng.2375 (2012).

32 Westenberger, S. J. et al. A systems-based analysis of Plasmodium vivax lifecycle transcription from human to mosquito. PLoS Negl Trop Dis 4, e653, doi:10.1371/journal.pntd.0000653(2010).

33 Su, X. Z. et al. The large diverse gene family var encodes proteins involved in cytoadherence and antigenic variation of Plasmodium falciparum-infected erythrocytes. Cell 82, 89–100 (1995).

34 Otto, T. D. et al. A comprehensive evaluation of rodent malaria parasite genomes and gene expression. BMC Biol 12, 86, doi:10.1186/s12915-014-0086-0 (2014).

35 Winter, G. et al. SURFIN is a polymorphic antigen expressed on Plasmodium falciparum merozoites and infected erythrocytes. J Exp Med 201, 1853–1863, doi:10.1084/jem.20041392 (2005).

36 Hiller, N. L. et al. A host-targeting signal in virulence proteins reveals a secretome in malarial infection. Science 306, 1934–1937, doi:10.1126/science.1102737(2004).

37 Crosnier, C. et al. Basigin is a receptor essential for erythrocyte invasion by Plasmodium falciparum. Nature 480, 534–537, doi:10.1038/nature10606 (2011).

38 Gruszczyk, J. et al. Structurally conserved erythrocyte-binding domain in Plasmodium provides a versatile scaffold for alternate receptor engagement. Proc Natl Acad Sci USA 113, E191–200, doi:10.1073/pnas.1516512113 (2016).

39 Iyer, J., Gruner, A. C., Renia, L., Snounou, G. & Preiser, P. R. Invasion of host cells by malaria parasites: a tale of two protein families. Mol Microbiol 65, 231–249, doi:10.1111/j.1365-2958.2007.05791.x (2007).

40 Menard, D. et al. Whole genome sequencing of field isolates reveals a common duplication of the Duffy binding protein gene in Malagasy Plasmodium vivax strains. PLoS Negl Trop Dis 7, e2489, doi:10.1371/journal.pntd.0002489(2013).

41 McKenna, A. et al. The Genome Analysis Toolkit: a MapReduce framework for analyzing next-generation DNA sequencing data. Genome Res 20, 1297–1303, doi:10.1101/gr.107524.110(2010).

42 Volkman, S. K. et al. A genome-wide map of diversity in Plasmodium falciparum. Nat Genet 39,113–119, doi:10.1038/ng1930 (2007).

43 Neafsey, D. E. et al. The malaria parasite Plasmodium vivax exhibits greater genetic diversity than Plasmodium falciparum. Nat Genet 44, 1046–1050, doi:10.1038/ng.2373 (2012).

44 Innan, H. Modified Hudson-Kreitman-Aguade test and two-dimensional evaluation of neutrality tests. Genetics 173, 1725–1733, doi:10.1534/genetics.106.056242(2006).

45 Nekrutenko, A., Makova, K. D. & Li, W.H. The K(A)/K(S) ratio test for assessing the protein-coding potential of genomic regions: an empirical and simulation study. Genome Res 12,198–202, doi:10.1101/gr.200901 (2002).

46 Kreitman, M. & Hudson, R. R. Inferring the evolutionary histories of the Adh and Adh-dup loci in Drosophila melanogaster from patterns of polymorphism and divergence. Genetics 127, 565–582 (1991).

47 Otto, T.D et al. Genome sequencing of chimpanzee malaria parasites reveals possible pathways of adaptation to human hosts. Nat Commun 5, 4754, doi:10.1038/ncomms5754 (2014).

48 Li, W., Boswell, R. & Wood, W. B. mag-1, a homolog of Drosophila mago nashi, regulates hermaphrodite germ-line sex determination in Caenorhabditis elegans.Dev Biol 218,172–182, doi:10.1006/dbio.1999.9593 (2000).

49 Edgar, R. C. MUSCLE: multiple sequence alignment with high accuracy and high throughput. Nucleic Acids Res 32, 1792–1797, doi:10.1093/nar/gkh340 (2004).

50 Auburn, S. et al. An effective method to purify Plasmodium falciparum DNA directly from clinical blood samples for whole genome high-throughput sequencing. PLoS One 6, e22213, doi:10.1371/journal.pone.0022213 (2011).

51 Ollomo, B. et al. A new malaria agent in African hominids. PLoS Pathog 5, e1000446, doi:10.1371/journal.ppat.1000446 (2009).

52 Oyola, S. O. et al. Optimized whole-genome amplification strategy for extremely AT-biased template. DNA Res 21, 661–671, doi:10.1093/dnares/dsu028(2014).

53 Kamau, E. et al. K13-propeller polymorphisms in Plasmodium falciparum parasites from sub-Saharan Africa. J Infect Dis 211, 1352–1355, doi:10.1093/infdis/jiu608(2015).

54 Bronner, I. F., Quail, M. A., Turner, D. J. & Swerdlow, H. Improved Protocols for Illumina Sequencing. Curr Protoc Hum Genet 80, 18 12 11–42, doi:10.1002/0471142905.hg1802s80(2014).

55 Chin, C. S. et al. Nonhybrid, finished microbial genome assemblies from long-read SMRT sequencing data. Nat Methods 10, 563–569, doi:10.1038/nmeth.2474 (2013).

56 Otto, T. D., Sanders, M., Berriman, M. & Newbold, C. Iterative Correction of Reference Nucleotides (iCORN) using second generation sequencing technology. Bioinformatics 26, 1704–1707, doi:10.1093/bioinformatics/btq269(2010).

57 Altschul, S. F. et al. Gapped BLAST and PSI-BLAST: a new generation of protein database search programs. Nucleic Acids Res 25, 3389–3402 (1997).

58 Zimin, A. V. et al. The MaSuRCA genome assembler. Bioinformatics 29, 2669–2677, doi:10.1093/bioinformatics/btt476 (2013).

59 Boetzer, M., Henkel, C. V., Jansen, H. J., Butler, D. & Pirovano, W. Scaffolding pre-assembled contigs using SSPACE. Bioinformatics 27, 578–579, doi:10.1093/bioinformatics/btq683(2011).

60 Nadalin, F., Vezzi, F. & Policriti, A. GapFiller: a de novo assembly approach to fill the gap within paired reads. BMC Bioinformatics 13 Suppl 14, S8, doi:10.1186/1471-2105-13-S14-S8(2012).

61 Tsai, I. J., Otto, T. D. & Berriman, M. Improving draft assemblies by iterative mapping and assembly of short reads to eliminate gaps. Genome Biol 11, R41, doi:10.1186/gb-2010-11-4-r41(2010).

62 Assefa, S., Keane, T. M., Otto, T. D., Newbold, C. & Berriman, M. ABACAS: algorithm-based automatic contiguation of assembled sequences. Bioinformatics 25,1968–1969, doi:10.1093/bioinformatics/btp347 (2009).

63 Otto, T. D., Dillon, G. P., Degrave, W. S. & Berriman, M. RATT: Rapid Annotation Transfer Tool. Nucleic Acids Res 39, e57, doi:10.1093/nar/gkq1268 (2011).

64 Stanke, M. et al. AUGUSTUS: ab initio prediction of alternative transcripts. Nucleic Acids Res 34, W435–439, doi:10.1093/nar/gkl200 (2006).

65 Nawrocki, E. P. et al. Rfam 12.0: updates to the RNA families database. Nucleic Acids Res 43, D130–137, doi:10.1093/nar/gku1063 (2015).

66 Rutherford, K. et al. Artemis: sequence visualization and annotation. Bioinformatics 16, 944–945 (2000).

67 Carver, T. J. et al. ACT: the Artemis Comparison Tool. Bioinformatics 21, 3422–3423, doi:10.1093/bioinformatics/bti553 (2005).

68 Li, L., Stoeckert, C. J., Jr. & Roos, D. S. OrthoMCL: identification of ortholog groups for eukaryotic genomes. Genome Res 13, 2178–2189, doi:10.1101/gr.1224503 (2003).

69 Talavera, G. & Castresana, J. Improvement of phylogenies after removing divergent and ambiguously aligned blocks from protein sequence alignments. Syst Biol 56, 564–577, doi:10.1080/10635150701472164 (2007).

70 Stamatakis, A., Ludwig, T. & Meier, H. RAxML-III: a fast program for maximum likelihood-based inference of large phylogenetic trees. Bioinformatics 21, 456–463, doi:10.1093/bioinformatics/bti191 (2005).

71 Stamatakis, A., Hoover, P. & Rougemont, J. A rapid bootstrap algorithm for the RAxML Web servers. Syst Biol 57, 758–771, doi:10.1080/10635150802429642(2008).

72 Lartillot, N., Lepage, T. & Blanquart, S. PhyloBayes 3: a Bayesian software package for phylogenetic reconstruction and molecular dating. Bioinformatics 25, 2286–2288, doi:10.1093/bioinformatics/btp368 (2009).

73 Guindon, S. et al. New algorithms and methods to estimate maximum-likelihood phylogenies: assessing the performance of PhyML 3.0. Syst Biol 59, 307–321, doi:10.1093/sysbio/syq010 (2010).

74 Li, H. et al. The Sequence Alignment/Map format and SAMtools. Bioinformatics 25, 2078–2079, doi:10.1093/bioinformatics/btp352 (2009).

75 Morgulis, A., Gertz, E. M., Schaffer, A. A. & Agarwala, R. A fast and symmetric DUST implementation to mask low-complexity DNA sequences. J Comput Biol 13,1028–1040, doi:10.1089/cmb.2006.13.1028 (2006).

76 Bradley, R. K. et al. Fast statistical alignment. PLoS Comput Biol 5, e1000392, doi:10.1371/journal.pcbi.1000392(2009).

77 Claessens, A. et al. Generation of antigenic diversity in Plasmodium falciparum by structured rearrangement of Var genes during mitosis. PLoS Genet 10, e1004812, doi:10.1371/journal.pgen.1004812 (2014).

78 Daniels, R. F. et al. Modeling malaria genomics reveals transmission decline and rebound in Senegal. Proc Natl Acad Sci U S A 112, 7067–7072, doi:10.1073/pnas.1505691112(2015).

79 Zhang, Y. I-TASSER server for protein 3D structure prediction. BMC Bioinformatics 9, 40, doi:10.1186/1471-2105-9-40 (2008).

80 Yang, J. et al. The I-TASSER Suite: protein structure and function prediction. Nat Methods 12, 7–8, doi:10.1038/nmeth.3213 (2015).

81 Zhang, Y. & Skolnick, J. TM-align: a protein structure alignment algorithm based on the TM-score. Nucleic Acids Res 33, 2302–2309, doi:10.1093/nar/gki524 (2005).

82 Bastian M., H. S., & Jacomy M.. Gephi: an open source software for exporing and manipulating networks. International AAAI Conference on Weblogs and Social Media(2009).

83 Fruchterman, T. M. J., & Reingold, E. M. Graph drawing by force-directed placement. Software: Practice and Experience 21,1129–1164 (1991).

84 Enright, A. J., Van Dongen, S. & Ouzounis, C. A. An efficient algorithm for large-scale detection of protein families. Nucleic Acids Res 30,1575–1584 (2002).

85 Ochoa, D. & Pazos, F. Studying the co-evolution of protein families with the Mirrortree web server. Bioinformatics 26, 1370–1371, doi:10.1093/bioinformatics/btq137(2010).

86 Holloway, A. K., Lawniczak, M. K., Mezey, J. G., Begun, D. J. & Jones, C. D. Adaptive gene expression divergence inferred from population genomics. PLoS Genet 3, 2007–2013, doi:10.1371/journal.pgen.0030187 (2007).

87 Nei, M. & Gojobori, T. Simple methods for estimating the numbers of synonymous and nonsynonymous nucleotide substitutions. Mol Biol Evol 3, 418–426 (1986).

88 Langmead, B. & Salzberg, S. L. Fast gapped-read alignment with Bowtie 2. Nat Methods 9, 357–359, doi:10.1038/nmeth.1923 (2012).

89 Roberts, A. & Pachter, L. Streaming fragment assignment for real-time analysis of sequencing experiments. Nat Methods 10, 71–73, doi:10.1038/nmeth.2251 (2013).

90 Chimpanzee, S. & Analysis, C. Initial sequence of the chimpanzee genome and comparison with the human genome. Nature 437, 69–87, doi:10.1038/nature04072 (2005).

91 Marmoset Genome, S. & Analysis, C. The common marmoset genome provides insight into primate biology and evolution. Nat Genet 46, 850–857, doi:10.1038/ng.3042 (2014).

92 de Arruda, M., Nardin, E. H., Nussenzweig, R. S. & Cochrane, A. H. Sero-epidemiological studies of malaria in Indian tribes and monkeys of the Amazon Basin of Brazil. Am J Trop Med Hyg 41, 379–385 (1989).

